# Uncertainty in blacklisting potential Pacific plant invaders using species distribution models

**DOI:** 10.1101/2024.12.11.627501

**Authors:** Valén Holle, Anna Rönnfeldt, Katrin Schifferle, Juliano Sarmento Cabral, Dylan Craven, Tiffany Knight, Hanno Seebens, Patrick Weigelt, Damaris Zurell

## Abstract

1. Invasive alien species pose a growing threat to global biodiversity, underscoring the need for evidence-based prevention strategies. Species distribution models (SDMs) are a widely used tool to estimate the potential distribution of alien species and to inform blacklists based on establishment risk. Yet, data limitations and modelling decisions can introduce uncertainty in these predictions. Here, we aim to quantify the contribution of four key sources of uncertainty in SDM-based blacklists: species occurrence data, environmental predictors, SDM algorithms, and thresholding methods for binarising predictions.
2. Focusing on 82 of the most invasive plant species on the Hawaiian Islands, we built SDMs to quantify their establishment potential in the Pacific region. To assess uncertainty, we systematically varied four modelling components: species occurrence data (native vs. global), environmental predictors (climatic vs. edapho-climatic), four SDM algorithms, and three thresholding methods. From these models, we derived blacklists using three alternative blacklisting definitions and quantified the variance in establishment risk scores and resulting species rankings attributable to each source of uncertainty.
3. SDMs showed fair predictive performance overall. Among the sources of uncertainty, thresholding method had the strongest and most consistent influence on risk scores across all three blacklist definitions but resulted in only minor changes in blacklist rankings. In contrast, algorithm choice had the most pronounced effect on blacklist rankings, followed by smaller but important effects of species occurrence data and environmental predictors. Notably, models based only on native occurrences often underestimated establishment potential.
4. SDMs can provide valuable support for planning the preventive management of alien species. However, our findings show that blacklist outcomes are highly sensitive to modelling decisions. While ensemble modelling across multiple algorithms is a recommended best practice, our results reinforce the importance of incorporating global occurrence data when available and carefully evaluating the trade-offs of including additional environmental predictors. Given the strong influence of thresholding on risk scores, we emphasise the need for transparent, context-specific threshold selection. More broadly, explicitly assessing uncertainty in SDM outputs can improve the robustness of blacklists and support scientifically informed, precautionary decision-making, particularly in data-limited situations where pragmatic modelling choices must be taken.

## Introduction

Invasive alien species have been identified as a major direct driver of native biodiversity loss on a global scale, and a threat to human health and ecosystem services (IPBES, 2023; Pyšek et al., 2020). The magnitude of these impacts is predicted to increase rapidly, as the breakdown of biogeographic barriers continues to be facilitated by the worldwide expansion of trade and transport (Seebens et al., 2021). To slow down the accumulation of alien species, evidence-based efforts in implementing effective management strategies, specifically targeting the prevention of biological invasions, are urgently needed (Seebens et al., 2017; Simberloff et al., 2013). In light of limited resources, such as financial and time constraints, priority-setting is key for planning adequate prevention measures and maximising expected conservation benefits (Bellard et al., 2017; Ziller et al., 2020). For this purpose, blacklists have proven to be an effective policy tool to regulate introductions by providing a ranked priority list of alien species that pose a high risk of becoming naturalised and invasive (Cerri et al., 2022; Ziller et al., 2020). Importantly, such blacklists should rely on objectively quantified invasion risks of species to serve as a robust decision tool (Gallien et al., 2019; Pili et al., 2024).

Correlative species distribution models (SDMs) have become a widely used decision-support tool for risk analysis (Li & Wang, 2013; Peterson et al., 2011). These models statistically relate observed species occurrences to current environmental conditions and can be used to predict a species’ potential distribution in non-native areas. SDMs are comparably easy to use, have low data requirements, and benefit from the increasing availability of global biodiversity and environmental data (Wüest et al., 2020). However, if SDMs are to inform management decisions, such as the creation of blacklists for invasive species, a thorough analysis of uncertainties related to methodological and fundamental limitations is essential (Barbet-Massin et al., 2018; Buisson et al., 2010; Sittaro et al., 2023). Previous work has shown that the choice of SDM algorithms can affect spatial and temporal predictions, especially under global change scenarios (Buisson et al., 2010; Thuiller et al., 2019). To mitigate this challenge and enhance the robustness of species distribution predictions, considering multiple algorithms in an ensemble modelling approach has become best practice (Araújo et al., 2019; Araujo & New, 2007). However, uncertainty in SDMs extends well beyond algorithm choice. Comparatively less attention has been paid to the sensitivity of SDMs to variation in species and environmental input data, which could be of particular relevance when modelling distributions of potentially invasive species (Beaumont et al., 2009). A further, often overlooked source of uncertainty is the thresholding method used to convert continuous occurrence probabilities into presence-absence predictions. Threshold choice can profoundly influence model performance, spatial predictions, and thus the estimated establishment risk of non-native species (Hellegers et al., 2025; Liu et al., 2013; Nenzén & Araújo, 2011). This has direct implications for conservation tools such as blacklists, where divergent thresholding criteria may result in inconsistent risk classifications. Taken together, these considerations underscore that a comprehensive evaluation of SDM uncertainty must go beyond algorithm choice to explicitly include the effects of input data (species occurrences and environmental predictors) and thresholding decisions. This is particularly important in applied contexts where SDMs guide regulatory decisions under data limitations and high stakes.

The choice of species input data is not trivial for alien species. SDMs are commonly perceived as modelling the realised ecological niche (Elith & Leathwick, 2009). However, while a species’ realised niche reflects the environmental (abiotic and biotic) conditions within its geographical range of current occurrences, evidence suggests that different realised niches could emerge when a species is introduced to a new geographical area (Arlé et al., 2024; Guisan et al., 2014). Possible reasons are associated with changes in the abiotic and biotic conditions promoting release from biotic constraints, changes in biotic interactions, or rapid genetic adaptation (Gallien et al., 2010; Guisan et al., 2014). These differences between native and non-native regions can ultimately lead to distinct realised niches, violating the underlying key assumption of niche conservatism in SDMs and their applications (Barbet-Massin et al., 2018). The reduced model transferability when extrapolating into new regions can therefore result in an underestimation of the species’ establishment potential (Beaumont et al., 2009). To address this issue, it has been suggested to include all available species occurrences from native and non-native areas together, to provide a more comprehensive estimation of the realised niche (Beaumont et al., 2009; Gallien et al., 2012; Liu et al., 2022). However, the use of such global data depends on the availability of sufficient non-native occurrence records, which are often lacking, especially in early-stage invasions or in data-poor regions. This trade-off between data completeness and niche coverage is a critical source of uncertainty affecting model predictions for potentially invasive species and their applicability for risk assessment.

Another important source of uncertainty lies in the selection of environmental predictors. These variables should capture the ecological requirements of a species to produce ecologically meaningful and accurate predictions (Descombes et al., 2020). Climate predictors are widely used to determine species distributions, presumably due to their recognised role as a primary determinant of species distributions and the high availability of global climate data (Guisan et al., 2017). But particularly for plants, the complementary consideration of soil properties in SDMs, such as soil pH and nitrogen content, can significantly improve prediction quality (Dubuis et al., 2013), suggesting that these could also be of relevance for predicting non-native plant distributions (Lázaro-Lobo et al., 2021; Sittaro et al., 2023). Yet, soil data are often derived from interpolated global maps with variable spatial coverage and regional accuracy (Poggio et al., 2021), which introduces further uncertainty. This creates a trade-off between richer environmental information and consistent spatial coverage that can impact model transferability, particularly when extrapolating to new regions. Together, uncertainties arising from species occurrence data and environmental predictors are especially relevant for invasive species management, where decision-makers often must base their decisions on patchy, sparse, or regionally biased data. Understanding how these modelling choices (algorithm choice, threshold choice, species data, environmental data) affect the robustness of blacklist rankings is essential for generating scientifically informed, actionable risk assessments.

To better understand the uncertainties linked with SDMs and their implications for blacklisting potentially invasive species, the Pacific region represents an ideal study system. The Pacific Islands have rich biodiversity and high levels of endemism that are highly threatened by biological invasions. Compared to other regions of the world, the Pacific region demonstrates the steepest regional relationship between alien plant species and area, meaning that the fastest increase in alien plant species worldwide is found on the Pacific islands (van Kleunen et al., 2015). The Hawaiian Islands, in particular, are strongly invaded, with nearly as many alien as there are native plant species. However, at this time, a large majority of the Pacific alien plant species are still spatially restricted to a few island groups (Wohlwend et al., 2021), suggesting that the adverse effects of biological invasions across much of the Pacific can be tackled by implementing preventive measures such as blacklists. Additionally, most of the Pacific alien plant species have also naturalised elsewhere in the world, making them a particularly useful study system to test the effects of species input data vis-à-vis environmental data, algorithm choice, and threshold choice on prediction and blacklisting uncertainty.

Here, we aim to construct Pacific-wide blacklists based on SDMs while explicitly considering uncertainty introduced by different species input data, environmental input data, SDM algorithms, and threshold selection methods. We focus on 82 of the most invasive species present on the Hawaiian Islands according to the PIER database (Pacific Island Ecosystems at Risk; http://www.hear.org/pier/), which could potentially spread to other Pacific island groups. Our study objectives were threefold: 1) to quantify the relative importance of the uncertainty factors on the blacklist ranking, considering differences in species data (native vs. global occurrences), environmental data (climatic vs. edapho-climatic predictors), SDM algorithm (regression-based and machine-learning), and thresholding methods, 2) to analyse variation in uncertainty between three different blacklist definitions, and 3) to analyse resulting differences in predictions of unrealised colonisation potential across the Pacific region.

## Methods

The general workflow involves fitting SDMs on opportunistic species occurrence data that represent the native or global niches of these species, using different sets of environmental predictors, SDM algorithms, and thresholding methods. From the predicted habitat suitability maps, we derived blacklists using three approaches. Additionally, we assess unrealised colonisation potential within the Pacific region by comparing our predictions of potential distribution against island-level checklists of established non-native species. Following the ODMAP protocol (Zurell, Franklin, et al., 2020), we detail all data preparation and modelling steps in Supplementary Material C.

### Species and environmental data

#### Species data

We focused on an initial subset of the 122 most invasive plant species, occurring on at least one of the Hawaiian Islands, whose invasive status is recognised by the PIER database (http://www.hear.org/pier/). The well-established commercial ties with much of the Pacific suggest that these species might spread to other Pacific Islands. The initial subset was derived from Wohlwend et al. (2021), containing information on the presence of non-native species on the different Pacific island groups. This dataset encompassed 50 island groups, defined both by political and geographic borders. These groups included islands located from 40°N to 40°S, ranging in size from 1 km² to 66,141 km². The delineation of the studied Pacific region was primarily based on biogeographic factors, as specific island features, such as geographic isolation and size, contribute to a particular vulnerability to the non-native establishment (Reaser et al., 2007). Therefore, we did not consider large archipelagos such as New Zealand, Japan, the Philippines, Indonesia, and Papua New Guinea, nor small islands located near the American and Australian Pacific coasts, nor islands belonging to Japan and South East Asia west of Bonin/Palau. The presence of non-native species at the island group level, as documented by Wohlwend et al. (2021), also served as checklist data to compare our blacklist predictions to species occurrence at the island group level and quantify unrealised colonisation potential across the Pacific region. For training SDMs, we acquired spatially-explicit occurrence data and derived two different species datasets: one containing only native occurrences and another containing both native and introduced (i.e., global) occurrences. These data were obtained from two databases, GBIF (Global Biodiversity Information Facility; https://www.gbif.org/; https://doi.org/10.15468/dd.vhjyha) and BIEN (Botanical Information and Ecology Network; https://bien.nceas.ucsb.edu/bien/; Supplementary Material A lists the acknowledged herbarium data sources). We cleaned the coordinates by excluding occurrences with erroneous time stamps and coordinates (Zizka et al., 2019) and removed duplicate records within 1 km² cells for each species to obtain a target spatial resolution of 1 km.

To compare SDMs fitted with native or global species data, we determined the biogeographical status of plant occurrences as native or introduced. We therefore consulted the following sources: the World Checklist of Vascular Plants (WCVP; https://powo.science.kew.org) at the base resolution of tdwg level 3 (Brummitt, 2001), the Global Inventory of Floras and Traits (GIFT; Weigelt et al., 2020), and the Global Naturalized Alien Flora (GloNAF; Pyšek et al., 2017; van Kleunen et al., 2015). The final biogeographical status was determined by merging the assignment information of all three sources. In the simplest case, all available source information agreed on the native or introduced status of each occurrence. When sources provided conflicting status assignments but referred to areas of different sizes, we assigned the status from the source referring to the smaller area. If an occurrence with conflicting sources referred to the same area size, we classified it as having a contradictory status. Overall, our final status assignments classified species occurrences as native, introduced, contradictory, or unknown.

Because our resulting species occurrence data only contained presence data, but most SDM algorithms require background data to identify areas with unsuitable environmental conditions, we derived pseudo-absences or background data for all study species (Barbet-Massin et al., 2012). For this, we randomly selected background points within a buffer distance of 200 km from the presences while excluding the presence locations. We chose this buffer distance to account for the biogeographic dispersal limitations of species. Following previous recommendations (Barbet-Massin et al., 2012), we used a presence-background ratio of 1:10 for the regression-based SDM algorithms and 1:1 for machine-learning algorithms. Lastly, to avoid spatial autocorrelation, all presences and background data of the species were thinned using a 3 km threshold (Aiello-Lammens et al., 2015).

For the final species selection, we set a lower limit of 40 native occurrences per species to avoid overfitting for species with a low sampling effort (Breiner et al., 2015). An additional criterion tested for a minimum difference of 40 between native and global occurrences to ensure the consideration of two different niches in the species’ input data. Species that, contrary to the information from Wohlwend et al. (2021), showed native records on the Hawaiian Islands were also excluded. The species selection process resulted in a final list of 82 species that were considered in further analyses (Supplementary B Table S1).

#### Environmental data

Similar to the species data, we built two different predictor datasets, one containing only climatic data and another containing climatic and edaphic data. These environmental predictor variables were obtained at a resolution of 1 km, matching the species data. Climate predictor variables were acquired from CHELSA (Climatologies at High Resolution for the Earth Land Surface Areas; version 2.1; Karger et al., 2017, 2018), including 15 bioclimatic variables. We excluded bioclimatic variables that combine temperature and precipitation data to avoid artefacts leading to discontinuities in interpolated surfaces (Booth, 2022). A set of 14 edaphic variables was obtained from SoilGrids (https://www.soilgrids.org), containing soil physical and chemical properties as well as soil water contents at different pressure heads (version 2.0; Poggio et al., 2021). Soil property maps were downloaded for three standard depth intervals from the surface to a depth of 30 cm and were then averaged over these depths (Descombes et al., 2020). The spatial coverage of the edaphic data was lower than that of the climate data, which mostly affected the maps of predicted occurrence probabilities. The lower spatial coverage was mainly attributable to urban areas and water surfaces. For predictions, we only included island groups with a representative spatial data coverage of ≥ 50 % for all variables. The island group of Tuvalu was not considered in any predictions due to its small size relative to the 1 km resolution used. Hence, we made predictions based on purely climatic data for 49 Pacific island groups, and predictions incorporating both predictor variable types for 25 island groups (Fig. 1; Supplementary B Table S2).

**Figure 1:**
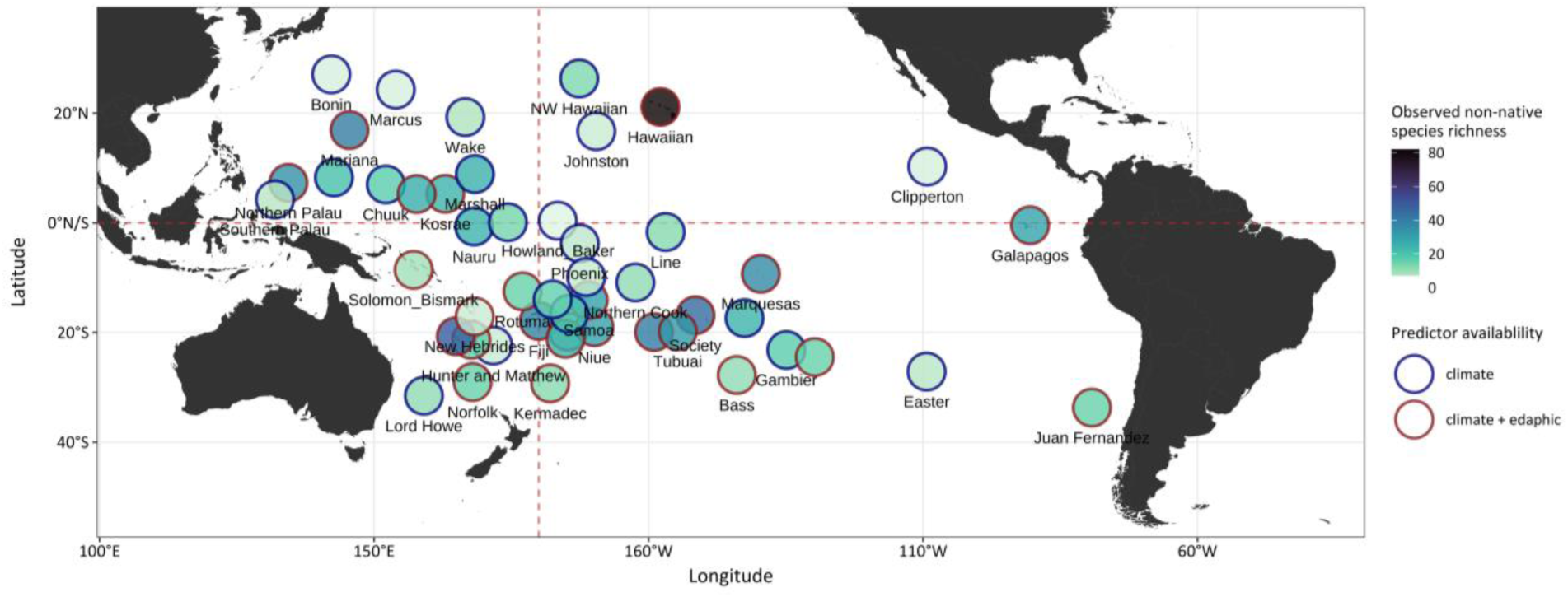
Study region. The location of Pacific island groups included in the study, each represented by a circle for easier visualisation. The circle shading shows the observed non-native species richness per island group, considering all studied non-native plant species (n = 82). Additionally, the availability of climatic and edaphic predictor variables per island group is indicated by the border colour, with a predictor type considered available if a representative spatial data coverage of ≥ 50 % per island group was achieved. Due to space limitations, not all island group labels are displayed. The circles are positioned at the central coordinates of each island group. The red dashed lines represent the equator (horizontal line) and the prime meridian (vertical line).

In accordance with our study objectives, the two types of species data (native vs. global occurrences) were intersected with the two types of environmental data (climatic vs. edapho-climatic predictors) resulting in four input datasets: ‘natclim’, which describes the combination of the native niche and purely climatic data, ‘natclim+eda’, which considers the native niche and edaphoclimatic data, and ‘globclim’ and ‘globclim+eda’, which describe the analogous input datasets regarding the global niche (Supplementary A Fig. S1).

### Species distribution models

For each of the four SDM input datasets per species, the four most important and weakly correlated predictor variables were identified prior to model building. To achieve this, we identified highly correlated pairs of variables with Spearman correlations |r|> 0.7 and only retained the most important variable (Dormann et al., 2013). Variable importance was determined using univariate generalised linear models (GLM) and a 5-fold cross-validation approach (Zurell, Zimmermann, et al., 2020). The univariate GLMs were fitted with a binomial error distribution and second-order polynomials, and with equal weight given to the sum of presences and background data. We then assessed the univariate predictive performance for each variable by cross-validation using the Boyce index (Hirzel et al., 2006). For SDMs that used both types of predictor variables, the two climatic and two edaphic variables that were ranked most important in the list of weakly correlated predictors were considered for model building (Supplementary B Table S3).

SDMs were fitted using four different statistical algorithms with intermediate model complexity: GLM, generalised additive models (GAM), random forest (RF) and boosted regression trees (BRT). GLMs were estimated with linear and quadratic terms, and GAMs were estimated with non-parametric cubic smoothing splines with up to four degrees of freedom. RF models were fitted with 1,000 trees and a minimum node size of 20. Finally, we fitted BRTs using a tree complexity of 2 and a bag fraction of 75 % and optimised the learning rate to obtain tree numbers between 1,000 and 5,000 (Elith et al., 2008). For the regression-based methods (GLM, GAM), we used all background data with a presence-background ratio of 1:10 and presences and background were equally weighted in model building. For the machine-learning methods (RF, BRT), we used ten replicate sets of background data in a ratio of 1:1 to the presences, resulting in 10 models each that were subsequently averaged (Barbet-Massin et al., 2012; Dullinger et al., 2017).

For all models, SDM performance was evaluated using a 5-fold cross-validation by computing the area under the receiver operating characteristic curve (AUC), true skill statistic (TSS), sensitivity, specificity, and the Boyce index. A model would be judged to have fair predictive performance if it achieved Boyce and AUC values > 0.7, and a TSS value > 0.5 (Dormann et al., 2013; Guisan et al., 2017). In line with our study objectives, three different thresholding methods were applied to derive threshold-dependent performance metrics (TSS, sensitivity, specificity). These included two thresholds proposed by Liu et al. (2013), which are less affected by background data: the threshold that maximises TSS (maxTSS) and the mean probability of training points (meanProb). Additionally, we employed a threshold commonly used in invasion biology, the 10th percentile of probabilities at training presences (tenthPer; Pearson et al., 2007). Cross-validated predictions for models based on regression methods used prior defined equal weights of the sum of presences and background, while for machine-learning algorithms, SDM performance in cross-validation was averaged over the ten replicate models (Barbet-Massin et al., 2012; Dullinger et al., 2017). Ensemble models were constructed using the arithmetic mean. The ensemble model performance was assessed based on the averaged cross-validated model predictions of all algorithms using the same input dataset and thresholding method. To guarantee equal contributions of the four considered model algorithms, the cross-validated predictions derived from the ten models of each machine-learning algorithm were averaged beforehand (Dullinger et al., 2017). Given that the maxTSS threshold is widely recognised as well suited for presence-background data (Liu et al., 2013), the main results presented hereafter refer to this threshold. Results obtained using the alternative thresholding methods are provided in Supplementary Material A.

### Blacklisting

We predicted occurrence probabilities and binary predictions of the potential presence and absence of all study species using the same procedure described above. Predicted presences were interpreted as indicating environmental conditions that define suitable habitats for the species. Based on these predictions we quantified establishment risk scores for each species and subsequently constructed blacklists to identify species with the highest potential for establishment across the Pacific. We used three different definitions for the risk scores and blacklists (Supplementary A Fig. S1): First, the total suitable habitat fraction across the Pacific region considered the establishment potential across the entire Pacific region independent of individual island group sizes. It was derived by summing up the predicted suitable area across island groups and dividing it by the total area of all island groups. Then, species were ranked according to their total predicted suitable habitat fraction, with the species with the highest habitat fraction receiving rank 1. Second, the blacklist based on the mean suitable habitat fraction across all Pacific island groups puts more emphasis on local establishment risk, highlighting the variability of establishment risk on different island groups. It was calculated by dividing the predicted suitable area per island group by the area of that island group and then averaging the suitable area fractions across all island groups. Species were then ranked according to their mean habitat fraction across all island groups, with the species with the highest mean fraction receiving rank 1. Third, the blacklist based on the number of island groups with suitable habitats does not consider habitat fraction per island group but simply uses a binary measure of whether suitable habitat is or is not found within an island group. An island group is considered to be potentially suitable if the fraction of potentially suitable habitat is greater than 0. Species were then ranked according to the number of island groups where they could establish. If species were predicted to have suitable habitats on the same number of island groups (regardless of their suitable habitat fraction on those island groups), they received the same rank. All blacklist definitions relied on the quantified absolute area of predicted suitable habitat in km² and the total area covered by data in km² at the island group level. For subsequent uncertainty analyses, we separately constructed a blacklist for each combination of species input data, environmental input data, SDM algorithm, and thresholding method. Additionally, we derived final blacklists with ensembles of SDM algorithms, to highlight how the choice of input data, threshold and blacklisting procedure would affect practical decision-making. To increase applied relevance, species ranks were further categorised into four distinct groups – Top 10, Top 20, Top 30, and Lower priority species – facilitating clearer interpretation of uncertainty-related impacts within a policy-relevant context.

Generally, the area of suitable habitat and the area of island groups were calculated based on spatial rasters with 1 km resolution. Thus, the calculated areas partly deviated from the true sizes of island groups, particularly those composed of numerous small-sized or thinly-shaped islands. Regarding the area of island groups, we considered the area covered by data instead of true island group sizes, as island groups with a data coverage of ≥ 50 % were included in SDM predictions. Due to differences in the spatial coverage of the environmental predictor datasets, resulting in differing numbers of considered island groups depending on the predictor type used, the ranking could only be compared when considering the same island groups. Thus, all final blacklist rankings were based on the 25 Pacific island groups covered by both predictor variable types (climatic and edaphic). Additionally, as a sensitivity analysis, we generated blacklists using purely climatic data considering prediction results for all 49 island groups covered by climatic data. This allowed for a more extensive analysis in terms of included island groups but did not allow us to assess environmental input data as a source of uncertainty in the resulting blacklists.

### Uncertainty analysis

We quantified four relevant sources of uncertainty in blacklisting potentially invasive species using SDMs: the species input data (native vs. global occurrences), the environmental input data (climatic vs. edapho-climatic predictors), SDM algorithms (GLM, GAM, RF, BRT), and thresholding methods (maxTSS, meanProb, tenthPer). Firstly, to quantify the relative importance of these uncertainties for blacklisting, we used a random forest approach to relate the establishment risk scores that serve for blacklist rankings, to categorical variables representing the choice of input data, algorithm, and threshold (Liaw & Wiener, 2002). Sources of uncertainty were quantified separately for each blacklist definition. Finally, we estimated the importance of each uncertainty factor, measured as the mean decrease in model accuracy when randomising the variables, which we normalised to add up to 100 %. Secondly, we directly assessed the impact of each uncertainty source on the actual blacklist rankings. For this, we calculated the mean difference in species rankings attributable to each factor across the three blacklist definitions. To provide a policy-relevant interpretation, species rankings were grouped into priority bins of ten, and changes in rank groupings were quantified as the proportion of cases exhibiting either stability or shifts within these bins.

### Unrealised colonisation potential

Finally, we quantified the potential for the species to establish on island groups beyond their current non-native range and how these predictions varied across input datasets and thresholding methods. For this, we calculated the Pacific-wide predicted unrealised colonisation potential per species, input dataset, and thresholding method using ensembles of SDM algorithms. Colonisation potential was estimated based on the island group-level checklists and defined as the False Positive Rate per island group (FPR = FP/(FP+TN)), which compares how many island groups that are hitherto unoccupied by a species would provide suitable habitat for that species. Reported presences at island group levels were taken from the checklists gathered by Wohlwend et al. (2021) (Supplementary B Table S4). A predicted presence on an island group was considered if the suitable habitat fraction was greater than zero. If a non-native species was unreported on an island group (absence) in the checklist but the model predicted a presence, then this prediction was considered a False Positive (FP), whereas non-native species that were neither reported nor predicted to occur were classified as True Negative (TN). In a final step, the FPR over all island groups was calculated for each study species. FPR ranges between 0 and 1, with 1 indicating that all island groups are unoccupied although providing suitable habitat, and 0 implying that none of the unoccupied island groups is predicted to provide suitable habitat for the studied species. In our main analyses, we considered the 25 island groups for which climatic and edaphic data availability allowed a comprehensive uncertainty assessment, but also repeated the analyses for the 49 island groups for which only climatic data were available.

For all analyses and visualisations, R 4.3.1 (R Core Team, 2023) was used, along with the following packages: *BIEN* (Maitner, 2023), *CoordinateCleaner* (Zizka et al., 2019), *dismo* (Hijmans et al., 2023), *ecospat* (Broennimann et al., 2023), *gbm* (Greenwell et al., 2022), *mgcv* (Wood et al., 2016), *PresenceAbsence* (Freeman & Moisen, 2008), *randomForest* (Liaw & Wiener, 2002), *rgbif* (Chamberlain et al., 2025), and *terra* (Hijmans, 2023).

## Results

### Model performance

The predictive performance of the SDMs, considering both threshold-independent and threshold-dependent metrics based on the maxTSS threshold, varied considerably among the studied species, ranging from poor to excellent. However, on average, fair performance scores over all models were achieved (TSS, mean ± SD: 0.52 ± 0.12; Boyce, mean ± SD: 0.94 ± 0.08; AUC, mean ± SD: 0.82 ± 0.07; Fig. 2; Supplementary A Fig. S2).

**Figure 2:**
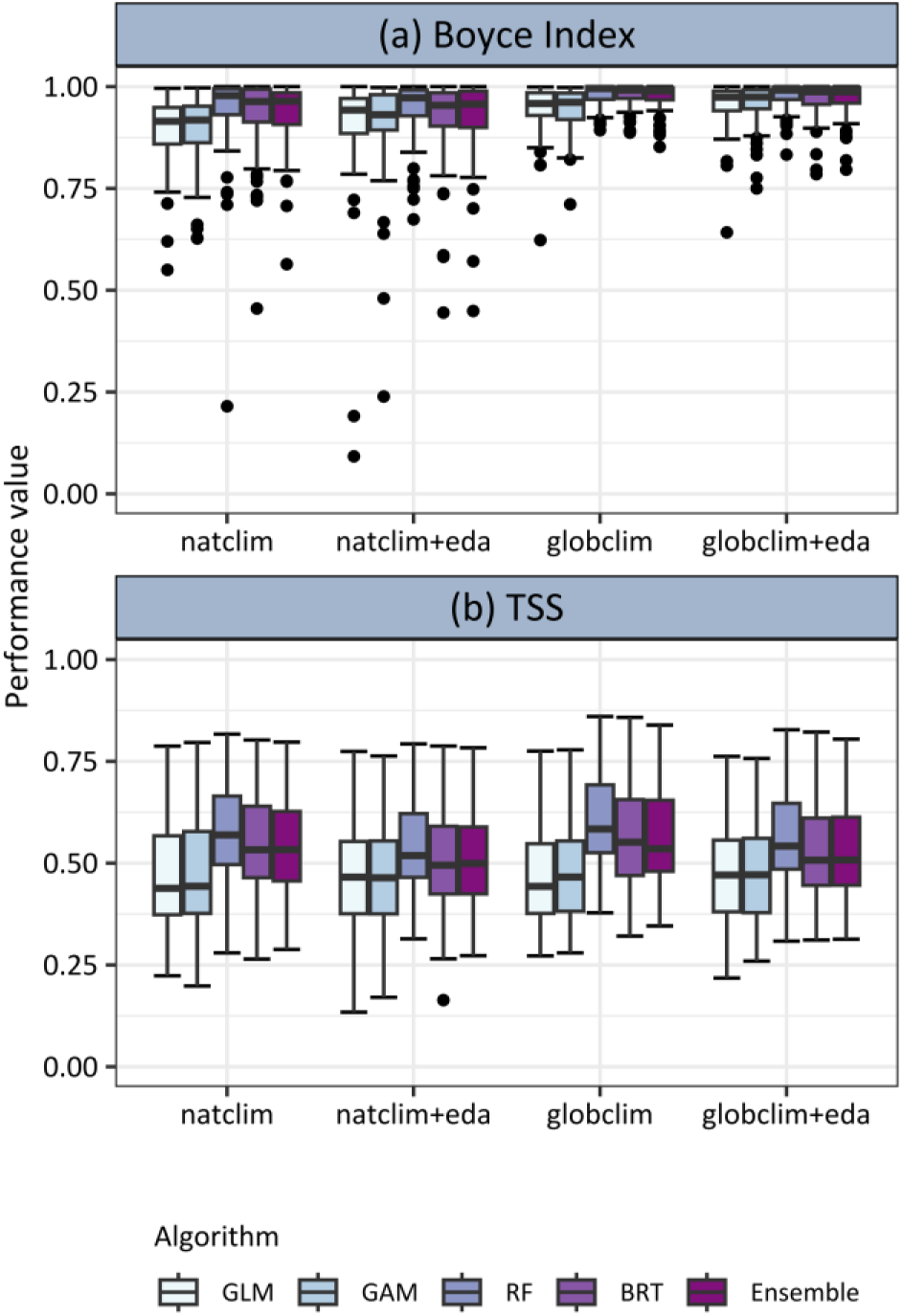
Model validation in terms of Boyce index and TSS (True Skill Statistic). For all studied non-native plant species (n = 82), the range of performance measures for the four applied model algorithms and the averaged ensemble model along the four different input datasets are displayed as boxplots (a-b). Models were validated using 5-fold cross-validation, and TSS values are shown based on the threshold that maximises the True Skill Statistic (maxTSS). (GLM: generalised linear model; GAM: generalised additive model; RF: random forest; BRT: boosted regression tree; natclim: models based on native occurrences and climate data; globclim: models based on global occurrences and climate data; eda: models additionally considering edaphic data)

Predictive performance also varied across SDM algorithms, with models based on the RF algorithm generally outperforming those based on the GLM, GAM, and BRT algorithms. This is demonstrated by an average TSS value of 0.57 ± 0.11 for all RF models, whereas average TSS scores of models based on the other algorithms were lower, ranging from 0.47 ± 0.12 to 0.54 ± 0.11 (Fig. 2b). Similarly, RF models achieved the highest average Boyce index (0.96 ± 0.07), although the differences were modest compared to other algorithms (GLM, mean ± SD: 0.92 ± 0.09; GAM, mean ± SD: 0.93 ± 0.08; BRT: mean ± SD: 0.95 ± 0.08; Fig. 2a). Ensemble models generally achieved fair performance accuracies (TSS, mean ± SD: 0.54 ± 0.11; Boyce, mean ± SD: 0.95 ± 0.07; AUC, mean ± SD: 0.84 ± 0.06; Fig. 2; Supplementary A Fig. S2).

Overall, the use of different input datasets resulted in smaller differences in predictive performance than the choice of SDM algorithm (Fig. 2; Supplementary A Fig. S2). Still, all models, including ensembles, achieved slightly higher model performances when trained on global rather than native occurrences. Also, the Boyce index showed markedly better model performances when considering global occurrences instead of native occurrences, with average values of 0.97 ± 0.05 and 0.91 ± 0.1, respectively (Fig. 2a).

When examining the threshold-dependent performance metrics of the alternative thresholding methods, meanProb and tenthPer consistently yielded lower average TSS values than maxTSS. While both thresholding methods achieved higher average sensitivity, this was accompanied by notably lower average specificity (Fig. 2; Supplementary A Fig. S2 + S3 + S4).

### Uncertainty analysis

Using the SDMs, we estimated establishment risk scores and constructed blacklists for 25 Pacific island groups and then quantified the influence of four key sources of uncertainty: species input data (native vs. global occurrences), environmental input data (climatic vs. edapho-climatic predictors), SDM algorithm, and thresholding method, across three blacklist definitions. We first evaluated how these sources affected the establishment risk scores underlying the blacklist rankings (Fig. 3). Among all factors, threshold choice emerged as the most influential source of uncertainty overall (explaining 32.5 % to 71.0 % of the variation in risk scores), although the relative importance of each factor varied depending on the blacklist definition. For the risk scores based on the number of suitable island groups, thresholding method (32.5 %), species occurrence data (34.7 %), and algorithm choice (32.8 %) contributed similarly and substantially to uncertainty. A similar pattern was observed for the mean suitable habitat fraction, although in this case, algorithm choice was comparably less influential (11.6 %) than thresholding (40.0 %) and species data (43.4 %). In contrast, for the blacklist based on the total suitable habitat fraction, thresholding was by far the most dominant contributor to uncertainty (71.0 %), followed by species occurrence data (29.0 %), whereas algorithm choice had no effect. Across all blacklist definitions, the selection of environmental predictors consistently had a low influence on establishment risk scores.

**Figure 3:**
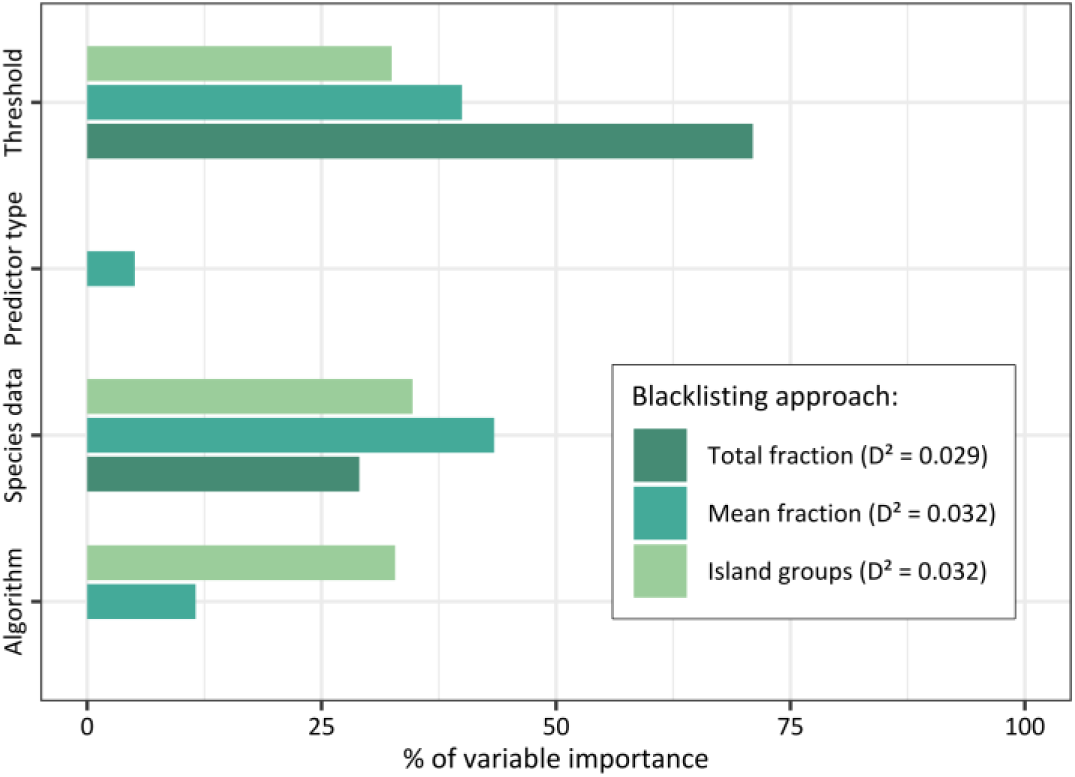
Quantification of uncertainty sources in establishment risk scores underlying blacklist rankings, based on analysis of 25 Pacific island groups. The bars display the influence in the choice of four different sources of uncertainty on variation in establishment risk scores: the SDM algorithm (GLM, GAM, RF, BRT), the species data (global vs. native occurrences), the environmental data (climatic vs. edapho-climatic predictors), and the thresholding method (maxTSS, meanProb, tenthPer). Variable importance was estimated using a random forest model that relates differences in establishment risk scores to the uncertainty factors. The values of variable importance per blacklisting approach were standardised to sum up to 100 %. The explained deviance D² of the models is noted in the legend. The models include establishment risk scores of all studied non-native plant species (n = 82) based on predictions to 25 Pacific island groups covered by climatic as well as edaphic data.

We then explored how these sources of uncertainty translated into differences in the blacklist rankings (Table 1). We found that a consistent pattern emerged across all three definitions: algorithm choice had the largest effect on species rankings, resulting in mean rank differences of up to 14.45 ranks and changes in grouping (e.g., between Top 10 or Top 20 ranks) in up to 80 % of cases. This was followed closely by the choice of species occurrence data and predictor type. Thresholding consistently caused the smallest ranking changes in both ranking positions and group assignments across all definitions.

**Table 1:** Quantification of uncertainty sources in blacklist rankings based on analysis of 25 island groups. For each blacklist approach, the table presents the mean differences in species rankings, along with the proportion of cases showing changes or no changes in rank groupings. Ranks were grouped into priority bins of 10 to assess grouping changes (i.e., Top 10, Top 20, Top 30, etc.). The analysis considers the four different uncertainty sources: the SDM algorithm (GLM, GAM, RF, BRT), the species data (global vs. native occurrences), the environmental data (climatic vs. edapho-climatic predictors), and the thresholding method (maxTSS, meanProb, tenthPer). Results are based on blacklisting assessments of all studied non-native plant species (n = 82), predicted across 25 Pacific island groups covered by climatic as well as edaphic data.

| Blacklist | Uncertainty source | Mean ranking differences | No change in grouping (%) | Change in grouping (%) |
| --- | --- | --- | --- | --- |
| Total fraction | Algorithm | 14.09 | 24 | 76 |
|  | Predictor type | 11.73 | 33 | 67 |
|  | Species data | 13.54 | 29 | 71 |
|  | Threshold | 7.84 | 41 | 59 |
| Mean fraction | Algorithm | 14.45 | 20 | 80 |
|  | Predictor type | 12.66 | 32 | 68 |
|  | Species data | 13.83 | 30 | 70 |
|  | Threshold | 7.75 | 43 | 57 |
| Island groups | Algorithm | 3.97 | 76 | 24 |
|  | Predictor type | 2.87 | 83 | 17 |
|  | Species data | 3.33 | 80 | 20 |
|  | Threshold | 2.28 | 86 | 14 |

The sensitivity analysis, based on suitability predictions from climate-only models for 49 island groups confirmed the same uncertainty patterns identified for the 25 Pacific island groups, both in terms of the establishment risk scores underlying the blacklist rankings and the rankings themselves (Supplementary A Fig. S5 + Table S1).

### Final blacklisting and unrealised colonisation potential

We provide final blacklist groupings based on ensemble predictions averaged across SDM algorithms and binarised using the maxTSS threshold (Fig. 4). Overall, a core subset of species consistently falls within high-risk categories (Top 10 – 30) across all four predictor sets, demonstrating a strong and stable risk signal regardless of predictor or species data type. Nonetheless, several species transition in and out of the high-risk categories depending on the predictor set used. These shifts in grouping are particularly evident when comparing models based on global versus native data, with a greater number of species showing fluctuations in their blacklist classifications. Importantly, these shifts often reflect a decrease in ranking; species that rank within high-risk categories under global data frequently fall into the lower-priority grouping (i.e., outside of the Top 30) when modelled using native-range occurrences. The inclusion of edaphic predictors appears to modify rankings for a subset of species, in some cases resulting in higher ranks, especially when using native-range models.

**Figure 4:**
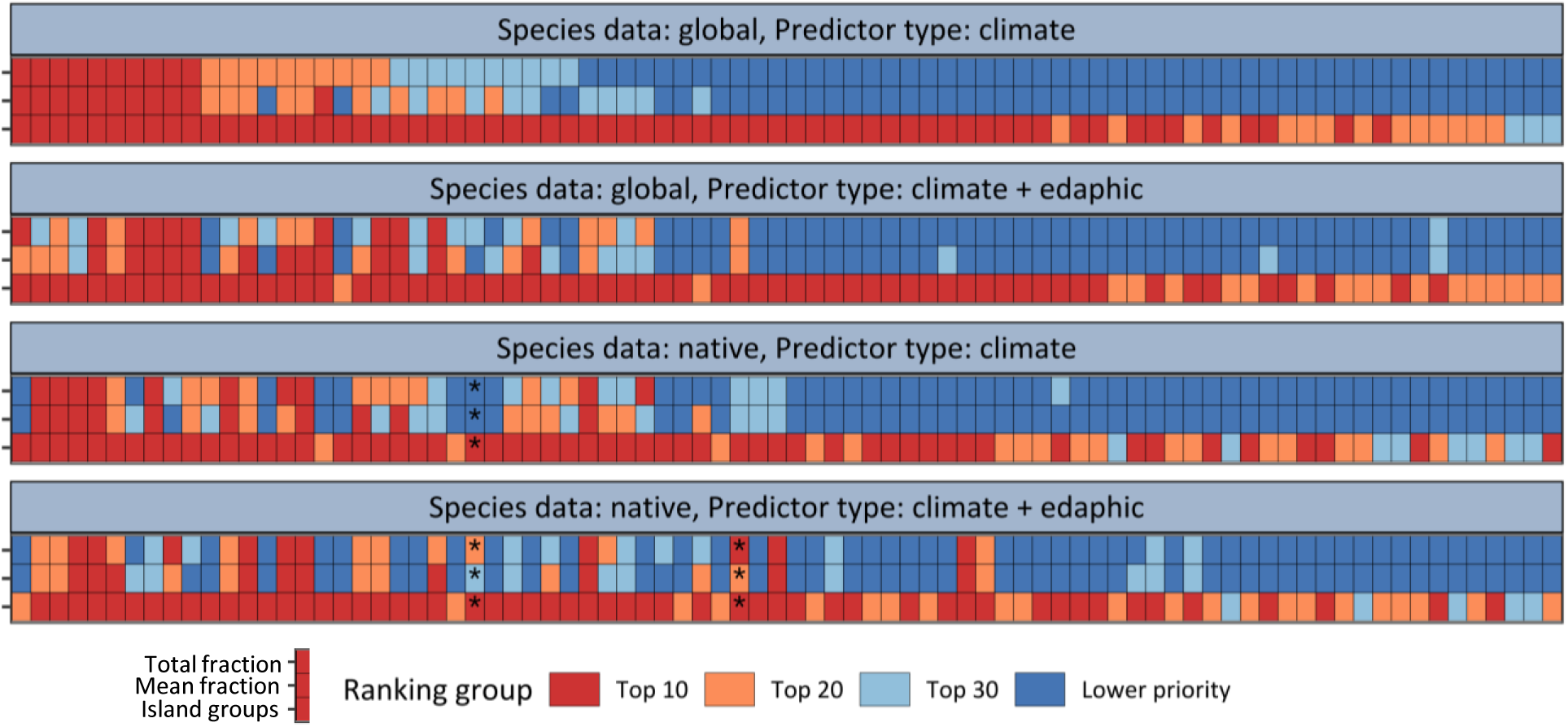
Final blacklist groupings based on analysis of 25 Pacific island groups. The final blacklist groupings of all three blacklisting approaches over the four different input datasets are displayed. The groupings are illustrated using the threshold that maximises the True Skill Statistic (maxTSS), based on the predictions of ensemble models to 25 Pacific island groups covered by climatic and edaphic data. Every non-native plant species (n = 82) is depicted along a vertical line of cells. The horizontal lines of cells per input dataset represent the three blacklisting approaches, with cell colour indicating the rank grouping of each species. Species ranks were categorised into four rank groups: Top 10, Top 20, Top 30, and Lower priority. Predictive performance values of ensemble models below or equal to a Boyce index of 0.7 are marked with an asterisk.

Focusing specifically on shifts towards lower-priority groupings, the non-native species *Echinochloa esculenta*, *Capsicum frutescens*, and *Lablab purpureus* exhibited the most pronounced changes, with rankings declining from the Top 10 to the Lower priority category under at least one of the blacklisting definitions. These shifts were directly attributable to variations in species data input from global to native occurrences. In addition, *Erigeron bonariensis* demonstrated an equally substantial shift in priority grouping, moving from the Top 10 to the Lower priority group when edaphic variables were not considered alongside climatic predictors.

Our sensitivity analysis, which examined climate-only models predicting habitat suitability for 49 Pacific island groups and incorporated two alternative thresholding approaches, revealed consistent patterns of ranking shifts across the four predictor sets (Supplementary A Fig. S6 + Fig. S7). Our final blacklists, derived from SDMs fitted with global occurrence data and edaphoclimatic predictors, and based on the maxTSS threshold, are provided for all three blacklisting definitions in Supplementary B Table S5.

From the ensemble predictions, we also assessed unrealised colonisation potential, i.e., the proportion of the 25 Pacific island groups with suitable habitats that have not yet been colonised by the current Hawaiian non-native species. We found that the choice of species data led to markedly different predictions. With an average of 0.66 ± 0.35, the unrealised colonisation potential was notably lower when only considering native occurrences. In comparison, when global occurrences were considered, the unrealised colonisation potential yielded an average value of 0.74 ± 0.34. The difference between occurrence datasets is largely due to the higher number of island groups that are predicted to harbour suitable habitat for Hawaiian non-native plant species (Fig. 5). This pattern persisted when predictions were extended to additional island groups using climate-only models, as well as under the alternative thresholding methods (Supplementary A Fig. S8 + S9).

**Figure 5:**
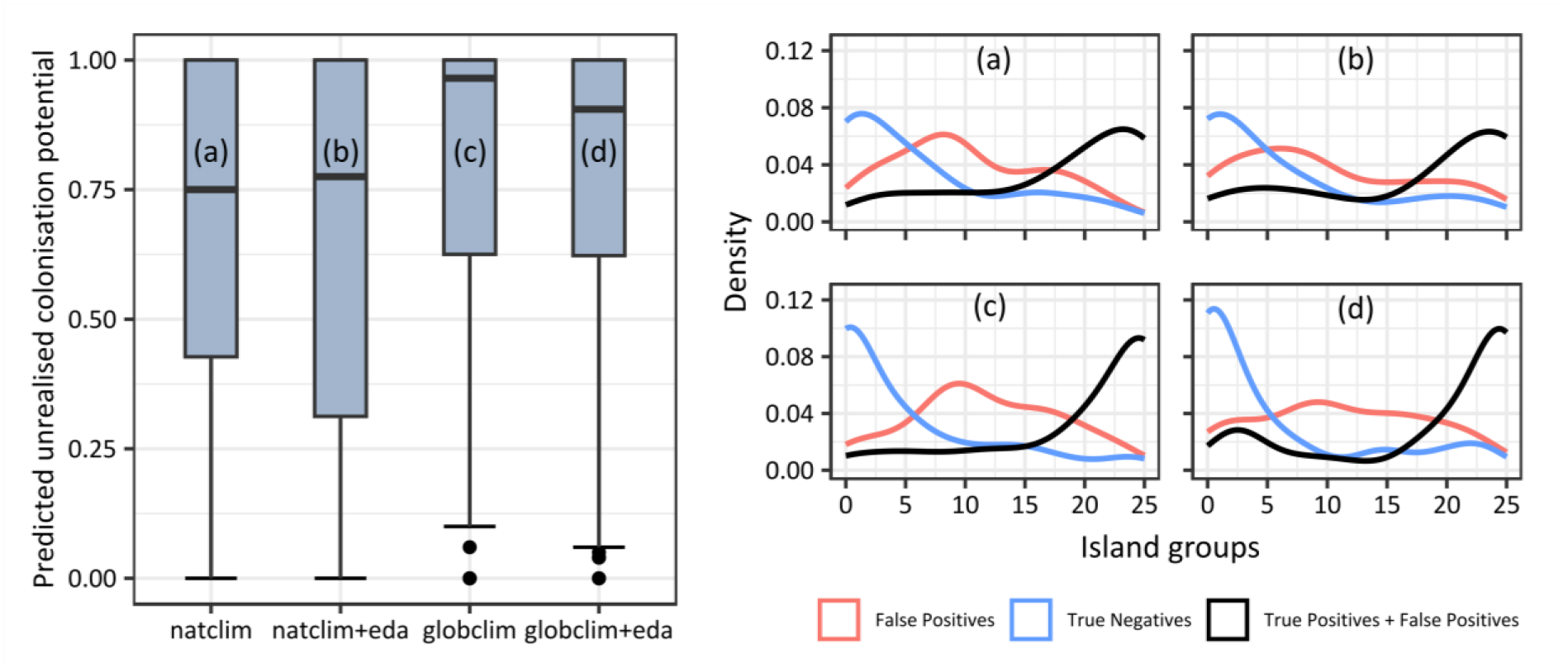
Predicted Pacific-wide unrealised colonisation potential based on analysis of 25 Pacific island groups. On the left, the predicted unrealised colonisation potential for all studied non-native plant species (n = 82) is displayed as boxplots, based on the ensemble predictions of 25 Pacific island groups covered by edaphic and climatic data along all four input datasets (a-d; natclim: models based on native occurrences and climate data; globclim: models based on global occurrences and climate data; eda: models additionally considering edaphic data). The predicted unrealised colonisation potential per species represents the False Positive Rate defined as the number of island groups that are hitherto unoccupied by the non-native species but predicted as suitable, divided by the total number of island groups where the species is currently absent. On the right, the density of island groups with false positives (predicted but absent; red line), true negatives (blue line), and all predicted presences (black line) is depicted for all study species using the same ensemble predictions (a-d). The predictions were binarised using the threshold that maximises the True Skill Statistic (maxTSS).

## Discussion

In this study, we quantified the contribution of four key sources of uncertainty in SDM-based blacklists by studying 82 of the most invasive plant species occurring on the Hawaiian Islands and assessing their establishment potential across the Pacific region. Our insights provide practical guidance for generating more robust and precautionary risk assessments, particularly in data-limited situations where managers must make critical decisions under uncertainty. Recognising that blacklists can be constructed in different ways and at different scales, we considered three different approaches to blacklisting potential invasive species by varying the emphasis on regional vs. local habitat suitability. Our results highlight the choice of thresholding method as the most consistent source of uncertainty across risk scores while the blacklist ranks varied most profoundly between different SDM algorithms, independent of the blacklist definition. Species data and environmental data had smaller, but non-negligible effects. Notably, when considering only native occurrences in SDMs, overall habitat suitability and thus unrealised colonisation potential was predicted to be lower across island groups, indicating that models based on native occurrences could provide overly optimistic risk scores and blacklists. These insights underscore the need for careful consideration of modelling choices in applied invasive species management and inform recommendations for improving the SDM-aided blacklisting in real-world decision contexts.

Our uncertainty analysis identified the choice of the SDM algorithm as the primary source of uncertainty in blacklist rankings. However, when focusing on the underlying establishment risk scores, the relative importance of algorithmic choice was lower and less consistent across blacklist definitions compared to other sources of uncertainty. To some extent, these findings reflect the known issue of variability in model outputs stemming from different algorithm choices. Multiple studies have reported such contrasting predictions arising from the use of different SDM algorithms, primarily in the context of climate change predictions (Buisson et al., 2010; Thuiller et al., 2019). Inconsistencies among models may stem from differences in model complexity and flexibility, particularly in the estimated species-environment relationships, which become even more pronounced when extrapolating into new niche spaces (Buisson et al., 2010). Additional to these known challenges, our results highlight a subtle but important aspect of SDM uncertainty in blacklisting: while algorithm choice strongly affected species rankings, its influence on absolute risk scores was less consistent. This likely reflects how ranking amplifies small differences in relative risk scores among species. Despite this sensitivity, ranking remains a necessary step in decision-making processes, as management actions often require prioritising species under limited resources. Understanding this duality highlights the importance of carefully considering algorithmic uncertainty when interpreting blacklist rankings to support scientifically informed management decisions.

The uncertainty stemming from species occurrence data substantially affected blacklisting across all three blacklisting approaches. This reflects the crucial influence that the availability of occurrence data has on the amount of predicted suitable habitat for non-native plant establishment, which in turn affects species rankings and predictions of unrealised colonisation potential. Models using global occurrences, as opposed to native occurrences, predicted larger areas of predicted suitable habitat and a higher number of suitable island groups across species. This suggests that some species may exhibit expansion of the realised niche between native and non-native regions, rather than conserving the realised niche from their native range (Guisan et al., 2014). Previous studies have shown such niche shifts and niche expansion to be common in invasive plant species (Barbosa et al., 2013; Beaumont et al., 2009; Gallagher et al., 2010). Invasive species have also been used as test cases for assessing the transferability of SDMs and accuracy in predicting ongoing invasions (Barbet-Massin et al., 2018; Jones, 2012). We highlighted some species for which the regional establishment potential was strongly underestimated when only considering native species occurrences; these were ranked as Lower priority based on native data, but were classified among the Top 10 on the blacklist when global occurrences were included. This could indicate niche expansion, e.g., due to competitive release. To verify this assumption, additional analyses would be necessary that specifically compare environmental niche overlap between native and non-native ranges (Gallien et al., 2012; Guisan et al., 2014). From a management perspective, relying solely on native occurrence data can limit model transferability and potentially overlook high-risk species. Yet, in many real-world situations, native occurrence data may be the main or only source of occurrence data, especially for early-stage invasions. Therefore, managers should flag these species for targeted monitoring and prioritise them for reassessment as new non-native occurrence data become available.

Among the four sources of uncertainty, the choice of environmental predictor type contributed comparably less to uncertainty in blacklisting outcomes and had only a minor influence on SDM performance. This suggests that climate predictors generally capture regional habitat suitability for the studied plant species. Previous studies found that including edaphic variables improved SDM predictive accuracy and thus play a crucial role in shaping plant distributions (Dubuis et al., 2013). However, these studies fitted SDMs at much finer spatial resolutions and focused on smaller spatial extents, underlining the importance of considering edaphic data when assessing plant habitat suitability more locally. Studies conducted at regional scales support our findings, highlighting climate as the overwhelming driver of plant species distributions (Beauregard & Blois, 2014; Syphard & Franklin, 2009). This demonstrates that small-scale variations of soil properties become less noticeable across broader regions or continents, while climate strongly determines the range margins of plant species distributions. Also, the lower importance of edaphic variables for the study of invasive plant species suggests that successful invaders may be generalists with respect to soil requirements. Nonetheless, we still observed non-negligible effects of environmental predictor choice on blacklist rankings. This indicates that although a less pronounced source of uncertainty, edaphic factors can still meaningfully alter blacklisting and prioritisation for certain species, especially those with more specific soil requirements. From a management perspective, this underlines the need for context-specific selection of predictor variables for modelling invasive plants. When soil data are available and of sufficient spatial coverage, their inclusion should be tested systematically, especially in local-scale risk assessments and for specialist plant species. However, global soil datasets often vary in spatial precision, interpolation accuracy and spatial coverage, especially on small islands and remote regions (Poggio et al., 2021), creating an important trade-off for model transferability. We thus recommend explicitly testing the effects of predictor types and data coverage on blacklisting outcomes to ensure reliability of resulting risk assessments.

The choice of thresholding method emerged as the most influential source of uncertainty when examining the establishment risk scores underlying blacklist rankings. However, its effect on the actual species rankings was comparably minor, suggesting a duality between thresholding effects on absolute risk estimates versus relative species prioritisation. This discrepancy indicates that while predicted risk scores can vary considerably depending on the thresholding approach, the relative species ranking tends to remain stable. This has important implications for invasive species management. Thresholding decisions become particularly critical in quantitative risk assessments, especially when comparing the magnitude of risk across regions or against predefined risk levels that may trigger policy action. In contrast, when managers must prioritise a fixed number of species for monitoring or intervention, the choice of threshold appears less consequential for relative species ranking. Despite this, thresholding decisions are rarely tested explicitly in SDM workflows (Hellegers et al., 2025; Nenzén & Araújo, 2011). Given their strong influence on risk quantification, we recommend that thresholding decisions be routinely tested and integrated into SDM workflows. Furthermore, thresholding decisions inherently affect the rate of false positives and false negatives in SDM-based risk assessments with potentially important consequences for invasive species management. In the case of false positives, we would erroneously blacklist non-native species with little chance of establishment, leading to unnecessary restrictions or resource allocation. In contrast, false negatives would mean that we are failing to blacklist non-native species with high establishment potential, which could pose ecological and economic risks through uncontrolled spread. The acceptable balance between these errors is highly context and resource-dependent. Transparently reporting the selected thresholding method and justifying it in light of the decision context will improve the interpretability of establishment risk and support clearer communication of those risks to stakeholders.

Our three alternative approaches to blacklisting potentially invasive plant species were designed to test the sensitivity of blacklists to habitat suitability with varying spatial emphasis. The ranking based on the total suitable habitat fraction reflects a broad, region-wide perspective, while the ranking based on the mean suitable habitat fraction takes a more local view, highlighting the variability of establishment risk on different island groups. The third blacklisting approach, which considers the number of island groups with suitable habitat, focuses on the presence or absence of suitable habitat, emphasising the potential reach of potentially invasive species regardless of the predicted extent of suitable habitat within each island group. These approaches are not exhaustive of potential blacklist definitions but nevertheless reflect a range of spatial objectives for blacklisting. Our results indicated that different species and environmental input data could lead to large discrepancies in ranking positions for all three blacklisting approaches. However, substantial ranking differences were attributable to only a few species, while the majority of blacklist rankings were robust. These findings underscore the importance of aligning blacklisting strategies with specific management goals. A broad regional risk assessment may prioritise different species than a local approach focused on protecting individual islands. Therefore, we recommend that the definition of blacklisting criteria should be carefully tailored to management objectives, their spatial scale and any additional constraints in available resources.

Our analyses show that many non-native species present on the Hawaiian Islands have a high colonisation potential across the Pacific island groups. Blacklists should prioritise species likely to establish that have also proven invasive in Hawaii and are not yet widespread in the Pacific. For instance, *Rhodomyrtus tomentosa*, a species native to southeastern Asia, is invasive in both Hawaii and Florida. Our results indicate that this species has a high potential to spread from the two island groups it currently occupies to all 25 considered island groups in the Pacific, making it a critical target for prevention (Smith et al., 2022). *Psidium guajava*, originally introduced as a fruit tree in Hawaii and many other regions, is now recognised as a major pest that aggressively displaces native species worldwide (Richardson & Rejmánek, 2011). In our study, *Psidium guajava* ranks high for all three blacklisting definitions. However, the window for effective prevention of this species may have passed, as it is already established on 22 island groups in the Pacific.

### Limitations and perspectives

We highlighted crucial factors contributing to uncertainties in blacklisting potentially invasive species based on SDMs. However, it is important to recognise that various aspects of SDMs can further contribute to uncertainty, as reflected by the low explanatory power of our uncertainty analysis. For instance, model parameter settings (Guisan et al., 2017) and sample size (Wisz et al., 2008) are additional factors that may affect model accuracy and subsequently lead to variation in spatial predictions. Moreover, potential biases associated with the underlying occurrence data could also have influenced the estimation of the species’ niches. To reduce such biases, we designed a number of data filtering steps and used several complementary data sources to build a robust framework for obtaining occurrence data and assigning their biogeographical status. Another factor influencing our results could be the limited availability of edaphic data, which substantially reduced our region of predictions (25 instead of 49 island groups). However, we are confident that our results are representative of the Pacific region, as our climate-only sensitivity analysis, which included almost twice as many island groups, yielded similar results. In the context of our study objectives, we assessed the establishment potential for all species as equivalent, based on the predicted suitable habitat identified by the fitted SDMs. If such information is available, blacklists could also consider additional factors relevant for establishment potential, for example, trait similarity with native flora (Denslow, 2003) or dominance capacity (Gallien et al., 2019). Another important consideration is the initial selection of species. Here, we focused our initial species selection on the PIER database and extracted all species listed as invasive on the Hawaiian Islands. However, depending on how often such databases are updated, blacklist applications intended to inform preventive measures may need to tap additional information sources to include the most current and comprehensive list of potential invasive species relevant to the region under study.

### Conclusions

While using ensembles of SDM algorithms is common and recommended for biodiversity assessments and decision-making (Araújo et al., 2019), our study demonstrates that other sources of uncertainty, such as species data, environmental predictors, and thresholding decisions, should be treated with equal scrutiny. Crucially, the relevance of each uncertainty source depends on the decision context and whether the aim is to communicate absolute risk levels, identify top-priority species, or inform a precautionary, resource-intensive management strategy versus an economically more efficient strategy.

We recommend that future blacklisting efforts should not only adopt ensemble modelling but also explicitly test the sensitivity of risk scores and blacklisting outcomes to input data and modelling choices. Embedding uncertainty analysis more routinely in SDM workflows will increase transparency and robustness of risk assessments and blacklists, helping managers to navigate trade-offs between precaution and feasibility, particularly in data-limited decision contexts.

## Author contributions

Valén Holle and Damaris Zurell conceived the ideas and designed the methodology; Valén Holle, Anna Rönnfeldt and Katrin Schifferle prepared the data; Valén Holle analysed the data; Valén Holle led the writing of the manuscript. All authors contributed critically to the drafts and gave final approval for publication.

## Acknowledgements

This study was supported by the German Research Foundation DFG (grant no. ZU 361/3-1 to DZ).

## Conflict of Interest

The authors declare that they have no conflict of interest to disclose.

## Data availability statement

The code used for the biogeographic status assignment can be accessed at https://github.com/UP-macroecology/StatusAssignment/releases/tag/v1.0.1, and the code for the uncertainty analyses at https://github.com/UP-macroecology/Holle_PacificPlantInvaders_BlacklistUncertainty_2023/releases/tag/v1.2.0.

## Notes

https://github.com/UP-macroecology/StatusAssignment/releases/tag/v1.0.1

https://github.com/UP-macroecology/Holle_PacificPlantInvaders_BlacklistUncertainty_2023/releases/tag/v1.2.0

### Competing Interest Statement

The authors have declared no competing interest.

### Summary of Updates

The revised version includes methodological updates and refinements aimed at highlighting the applied relevance of the results. Specifically, the following key updates were made: 1.Uncertainty analysis of thresholding methods: We extended the uncertainty analysis to explore the impact of different thresholding methods on blacklisting outcomes. Threshold selection is now explicitly treated as an additional source of uncertainty, enhancing the robustness of our modelling approach. 2.Refinement of uncertainty quantification: We expanded and deepened our uncertainty analysis to quantify the influence of each source of uncertainty not only on the blacklist rankings, but also on the underlying establishment risk scores that inform these rankings. This approach provides a more comprehensive understanding of how individual sources of uncertainty affect risk estimates. 3.Improved applied relevance: To increase the practical utility of our findings for practitioners, we reorganised the blacklist results into priority-bins (e.g., Top 10, Top 20, Top 30) as part of our uncertainty assessment and in the final blacklisting output. This format facilitates a clearer interpretation of results for end-users, such as policymakers, by identifying actionable priorities rather than presenting a continuous, and potentially less accessible, ranking. Additionally, we have revised relevant sections in the Abstract and Discussion to more explicitly articulate implications of uncertainty in SDM-based blacklists for invasive species management. In addition, some numerical values in the model performance and unrealised colonisation potential sections have been updated due to methodological refinements. These adjustments do not alter any conclusions or interpretations of the results.

## References

1. Aiello-Lammens, M. E., Boria, R. A., Radosavljevic, A., Vilela, B., & Anderson, R. P. (2015). spThin: An R package for spatial thinning of species occurrence records for use in ecological niche models. Ecography, 38(5), 541–545. 10.1111/ecog.01132

2. Araújo, M. B., Anderson, R. P., Márcia Barbosa, A., Beale, C. M., Dormann, C. F., Early, R., Garcia, R. A., Guisan, A., Maiorano, L., Naimi, B., O’Hara, R. B., Zimmermann, N. E., & Rahbek, C. (2019). Standards for distribution models in biodiversity assessments. Science Advances, 5(1), eaat4858. 10.1126/sciadv.aat4858

3. Araujo, M., & New, M. (2007). Ensemble forecasting of species distributions. Trends in Ecology & Evolution, 22(1), 42–47. 10.1016/j.tree.2006.09.010

4. Arlé, E., Knight, T. M., Jiménez-Muñoz, M., Biancolini, D., Belmaker, J., & Meyer, C. (2024). The cumulative niche approach: A framework to assess the performance of ecological niche model projections. Ecology and Evolution, 14(2), e11060. 10.1002/ece3.11060

5. Barbet-Massin, M., Jiguet, F., Albert, C. H., & Thuiller, W. (2012). Selecting pseudo-absences for species distribution models: How, where and how many? Methods in Ecology and Evolution, 3(2), 327–338. 10.1111/j.2041-210X.2011.00172.x

6. Barbet-Massin, M., Rome, Q., Villemant, C., & Courchamp, F. (2018). Can species distribution models really predict the expansion of invasive species? PLOS ONE, 13(3), e0193085. 10.1371/journal.pone.0193085

7. Barbosa, F. G., Pillar, V. D., Palmer, A. R., & Melo, A. S. (2013). Predicting the current distribution and potential spread of the exotic grass Eragrostis plana Nees in South America and identifying a bioclimatic niche shift during invasion. Austral Ecology, 38(3), 260–267. 10.1111/j.1442-9993.2012.02399.x

8. Beaumont, L. J., Gallagher, R. V., Thuiller, W., Downey, P. O., Leishman, M. R., & Hughes, L. (2009). Different climatic envelopes among invasive populations may lead to underestimations of current and future biological invasions. Diversity and Distributions, 15(3), 409–420. 10.1111/j.1472-4642.2008.00547.x

9. Beauregard, F., & Blois, S. de. (2014). Beyond a Climate-Centric View of Plant Distribution: Edaphic Variables Add Value to Distribution Models. PLOS ONE, 9(3), e92642. 10.1371/journal.pone.0092642

10. Bellard, C., Rysman, J.-F., Leroy, B., Claud, C., & Mace, G. M. (2017). A global picture of biological invasion threat on islands. Nature Ecology & Evolution, 1(12), 1862–1869. 10.1038/s41559-017-0365-6

11. Booth, T. H. (2022). Checking bioclimatic variables that combine temperature and precipitation data before their use in species distribution models. Austral Ecology, 47(7), 1506–1514. 10.1111/aec.13234

12. Breiner, F. T., Guisan, A., Bergamini, A., & Nobis, M. P. (2015). Overcoming limitations of modelling rare species by using ensembles of small models. Methods in Ecology and Evolution, 6(10), 1210–1218. 10.1111/2041-210X.12403

13. Broennimann, O., Cola, V. D., & Guisan, A. (2023). ecospat: Spatial Ecology Miscellaneous Methods. https://CRAN.R-project.org/package=ecospat

14. Brummitt, R. K. (with International Working Group on Taxonomic Databases for Plant Sciences & Hunt Institute for Botanical Documentation). (2001). World geographical scheme for recording plant distributions (ed. 2). Hunt Inst. for Botanical Documentation.

15. Buisson, L., Thuiller, W., Casajus, N., Lek, S., & Grenouillet, G. (2010). Uncertainty in ensemble forecasting of species distribution. Global Change Biology, 16(4), 1145–1157. 10.1111/j.1365-2486.2009.02000.x

16. Cerri, J., Carnevali, L., Monaco, A., Genovesi, P., & Bertolino, S. (2022). Blacklists do not necessarily make people curious about invasive alien species. A case study with Bayesian structural time series and Wikipedia searches about invasive mammals in Italy. NeoBiota, 71, 113–128. 10.3897/neobiota.71.69422

17. Chamberlain, S., Barve, V., Mcglinn, D., Oldoni, D., Desmet, P., Geffert, L., & Ram, K. (2025). rgbif: Interface to the Global Biodiversity Information Facility API. https://CRAN.R-project.org/package=rgbif

18. Denslow, J. S. (2003). Weeds in Paradise: Thoughts on the Invasibility of Tropical Islands. Annals of the Missouri Botanical Garden, 90(1), 119. 10.2307/3298531

19. Descombes, P., Walthert, L., Baltensweiler, A., Meuli, R. G., Karger, D. N., Ginzler, C., Zurell, D., & Zimmermann, N. E. (2020). Spatial modelling of ecological indicator values improves predictions of plant distributions in complex landscapes. Ecography, 43(10), 1448–1463. 10.1111/ecog.05117

20. Dormann, C. F., Elith, J., Bacher, S., Buchmann, C., Carl, G., Carré, G., Marquéz, J. R. G., Gruber, B., Lafourcade, B., Leitão, P. J., Münkemüller, T., McClean, C., Osborne, P. E., Reineking, B., Schröder, B., Skidmore, A. K., Zurell, D., & Lautenbach, S. (2013). Collinearity: A review of methods to deal with it and a simulation study evaluating their performance. Ecography, 36(1), 27–46. 10.1111/j.1600-0587.2012.07348.x

21. Dubuis, A., Giovanettina, S., Pellissier, L., Pottier, J., Vittoz, P., & Guisan, A. (2013). Improving the prediction of plant species distribution and community composition by adding edaphic to topo-climatic variables. Journal of Vegetation Science, 24(4), 593–606. 10.1111/jvs.12002

22. Dullinger, I., Wessely, J., Bossdorf, O., Dawson, W., Essl, F., Gattringer, A., Klonner, G., Kreft, H., Kuttner, M., Moser, D., Pergl, J., Pyšek, P., Thuiller, W., van Kleunen, M., Weigelt, P., Winter, M., Dullinger, S., & Beaumont, L. (2017). Climate change will increase the naturalization risk from garden plants in Europe. Global Ecology and Biogeography, 26(1), 43–53. 10.1111/geb.12512

23. Elith, J., & Leathwick, J. R. (2009). Species Distribution Models: Ecological Explanation and Prediction Across Space and Time. Annual Review of Ecology, Evolution, and Systematics, 40(1), 677–697. 10.1146/annurev.ecolsys.110308.120159

24. Elith, J., Leathwick, J. R., & Hastie, T. (2008). A working guide to boosted regression trees. Journal of Animal Ecology, 77(4), 802–813. 10.1111/j.1365-2656.2008.01390.x

25. Freeman, E. A., & Moisen, G. (2008). PresenceAbsence: An R Package for Presence Absence Analysis. Journal of Statistical Software, 23(11), 1–31. 10.18637/jss.v023.i11

26. Gallagher, R. V., Beaumont, L. J., Hughes, L., & Leishman, M. R. (2010). Evidence for climatic niche and biome shifts between native and novel ranges in plant species introduced to Australia. Journal of Ecology, 98(4), 790– 799. 10.1111/j.1365-2745.2010.01677.x

27. Gallien, L., Douzet, R., Pratte, S., Zimmermann, N. E., & Thuiller, W. (2012). Invasive species distribution models – how violating the equilibrium assumption can create new insights. Global Ecology and Biogeography, 21(11), 1126–1136. 10.1111/j.1466-8238.2012.00768.x

28. Gallien, L., Münkemüller, T., Albert, C. H., Boulangeat, I., & Thuiller, W. (2010). Predicting potential distributions of invasive species: Where to go from here? Diversity and Distributions, 16(3), 331–342. 10.1111/j.1472-4642.2010.00652.x

29. Gallien, L., Thornhill, A. H., Zurell, D., Miller, J. T., & Richardson, D. M. (2019). Global predictors of alien plant establishment success: Combining niche and trait proxies. Proceedings of the Royal Society B: Biological Sciences, 286(1897), 20182477. 10.1098/rspb.2018.2477

30. Greenwell, B., Boehmke, B., Cunningham, J., & Developers, G. B. M. (2022). gbm: Generalized Boosted Regression Models. https://CRAN.R-project.org/package=gbm

31. Guisan, A., Petitpierre, B., Broennimann, O., Daehler, C., & Kueffer, C. (2014). Unifying niche shift studies: Insights from biological invasions. Trends in Ecology & Evolution, 29(5), 260–269. 10.1016/j.tree.2014.02.009

32. Guisan, A., Thuiller, W., & Zimmermann, N. E. (2017). Habitat Suitability and Distribution Models: With Applications in R. Cambridge University Press. 10.1017/9781139028271

33. Hellegers, M., van Hinsberg, A., Lenoir, J., Dengler, J., Huijbregts, M. A. J., & Schipper, A. M. (2025). Multiple Threshold-Selection Methods Are Needed to Binarise Species Distribution Model Predictions. Diversity and Distributions, 31(4), e70019. 10.1111/ddi.70019

34. Hijmans, R. J. (2023). terra: Spatial Data Analysis. https://CRAN.R-project.org/package=terra

35. Hijmans, R. J., Phillips, S., Leathwick, J., & Elith, J. (2023). dismo: Species Distribution Modeling. https://CRAN.R-project.org/package=dismo

36. Hirzel, A. H., Le Lay, G., Helfer, V., Randin, C., & Guisan, A. (2006). Evaluating the ability of habitat suitability models to predict species presences. Ecological Modelling, 199(2), 142–152. 10.1016/j.ecolmodel.2006.05.017

37. IPBES. (2023). Thematic Assessment Report on Invasive Alien Species and their Control of the Intergovernmental Science-Policy Platform on Biodiversity and Ecosystem Services (H. E. Roy, A. Pauchard, P. Stoett, & T. Renard Truong, Eds.). IPBES Secretariat. 10.5281/zenodo.7430682

38. Jones, C. C. (2012). Challenges in predicting the future distributions of invasive plant species. Forest Ecology and Management, 284, 69–77. 10.1016/j.foreco.2012.07.024

39. Karger, D. N., Conrad, O., Böhner, J., Kawohl, T., Kreft, H., Soria-Auza, R. W., Zimmermann, N. E., Linder, H. P., & Kessler, M. (2017). Climatologies at high resolution for the earth’s land surface areas. Scientific Data, 4(1), Article 1. 10.1038/sdata.2017.122

40. Karger, D. N., Conrad, O., Böhner, J., Kawohl, T., Kreft, H., Soria-Auza, R. W., Zimmermann, N. E., Linder, H. P., & Kessler, M. (2018). Data from: Climatologies at high resolution for the earth’s land surface areas (Version 1, p. 7266970904 bytes) [Dataset]. Dryad. 10.5061/DRYAD.KD1D4

41. Lázaro-Lobo, A., Ramirez-Reyes, C., Lucardi, R. D., & Ervin, G. N. (2021). Multivariate analysis of invasive plant species distributions in southern US forests. Landscape Ecology, 36(12), 3539–3555. 10.1007/s10980-021-01326-3

42. Li, X., & Wang, Y. (2013). Applying various algorithms for species distribution modelling. Integrative Zoology, 8(2), 124–135. 10.1111/1749-4877.12000

43. Liaw, A., & Wiener, M. (2002). Classification and Regression by randomForest. R News, 2(3), 18–22.

44. Liu, C., White, M., & Newell, G. (2013). Selecting thresholds for the prediction of species occurrence with presence-only data. Journal of Biogeography, 40(4), 778–789. 10.1111/jbi.12058

45. Liu, C., Wolter, C., Courchamp, F., Roura-Pascual, N., & Jeschke, J. M. (2022). Biological invasions reveal how niche change affects the transferability of species distribution models. Ecology, 103(8), e3719. 10.1002/ecy.3719

46. Maitner, B. (2023). BIEN: Tools for Accessing the Botanical Information and Ecology Network Database. https://CRAN.R-project.org/package=BIEN

47. Nenzén, H. K., & Araújo, M. B. (2011). Choice of threshold alters projections of species range shifts under climate change. Ecological Modelling, 222(18), 3346–3354. 10.1016/j.ecolmodel.2011.07.011

48. Pearson, R. G., Raxworthy, C. J., Nakamura, M., & Townsend Peterson, A. (2007). ORIGINAL ARTICLE: Predicting species distributions from small numbers of occurrence records: a test case using cryptic geckos in Madagascar. Journal of Biogeography, 34(1), 102–117. 10.1111/j.1365-2699.2006.01594.x

49. Peterson, A. T., Soberón, J., Pearson, R. G., Anderson, R. P., Martínez-Meyer, E., Nakamura, M., & Araújo, M. B. (2011). Ecological Niches and Geographic Distributions (MPB-49). Princeton University Press. https://www.jstor.org/stable/j.ctt7stnh

50. Pili, A. N., Leroy, B., Measey, J. G., Farquhar, J. E., Toomes, A., Cassey, P., Chekunov, S., Grenié, M., van Winkel, D., Maria, L., Diesmos, M. L. L., Diesmos, A. C., Zurell, D., Courchamp, F., & Chapple, D. G. (2024). Forecasting potential invaders to prevent future biological invasions worldwide. Global Change Biology, 30(7), e17399. 10.1111/gcb.17399

51. Poggio, L., de Sousa, L. M., Batjes, N. H., Heuvelink, G. B. M., Kempen, B., Ribeiro, E., & Rossiter, D. (2021). SoilGrids 2.0: Producing soil information for the globe with quantified spatial uncertainty. SOIL, 7(1), 217–240. 10.5194/soil-7-217-2021

52. Pyšek, P., Hulme, P. E., Simberloff, D., Bacher, S., Blackburn, T. M., Carlton, J. T., Dawson, W., Essl, F., Foxcroft, L. C., Genovesi, P., Jeschke, J. M., Kühn, I., Liebhold, A. M., Mandrak, N. E., Meyerson, L. A., Pauchard, A., Pergl, J., Roy, H. E., Seebens, H., … Richardson, D. M. (2020). Scientists’ warning on invasive alien species. Biological Reviews, 95(6), 1511–1534. 10.1111/brv.12627

53. Pyšek, P., Pergl, J., Essl, F., Lenzner, B., Dawson, W., Kreft, H., Weigelt, P., Winter, M., Kartesz, J., Nishino, M., Antonova, L. A., Barcelona, J. F., Cabezas, F. J., Cárdenas, D., Cárdenas-Toro, J., Castaño, N., Chacón, E., Chatelain, C., Dullinger, S., … Kleunen, M. V. (2017). Naturalized alien flora of the world: Species diversity, taxonomic and phylogenetic patterns, geographic distribution and global hotspots of plant invasion. Preslia, 89(3), 203–274. 10.23855/preslia.2017.203

54. R Core Team. (2023). R: A Language and Environment for Statistical Computing. R Foundation for Statistical Computing. https://www.R-project.org/

55. Reaser, J. K., Meyerson, L. A., Cronk, Q., De Poorter, M., Eldrege, L. G., Green, E., Kairo, M., Latasi, P., Mack, R. N., Mauremootoo, J., O’Dowd, D., Orapa, W., Sastroutomo, S., Saunders, A., Shine, C., Thrainsson, S., & Vaiutu, L. (2007). Ecological and socioeconomic impacts of invasive alien species in island ecosystems. Environmental Conservation, 34(2), 98–111. 10.1017/S0376892907003815

56. Richardson, D. M., & Rejmánek, M. (2011). Trees and shrubs as invasive alien species – a global review. Diversity and Distributions, 17(5), 788–809. 10.1111/j.1472-4642.2011.00782.x

57. Seebens, H., Bacher, S., Blackburn, T. M., Capinha, C., Dawson, W., Dullinger, S., Genovesi, P., Hulme, P. E., van Kleunen, M., Kühn, I., Jeschke, J. M., Lenzner, B., Liebhold, A. M., Pattison, Z., Pergl, J., Pyšek, P., Winter, M., & Essl, F. (2021). Projecting the continental accumulation of alien species through to 2050. Global Change Biology, 27(5), 970–982. 10.1111/gcb.15333

58. Seebens, H., Blackburn, T. M., Dyer, E. E., Genovesi, P., Hulme, P. E., Jeschke, J. M., Pagad, S., Pyšek, P., Winter, M., Arianoutsou, M., Bacher, S., Blasius, B., Brundu, G., Capinha, C., Celesti-Grapow, L., Dawson, W., Dullinger, S., Fuentes, N., Jäger, H., … Essl, F. (2017). No saturation in the accumulation of alien species worldwide. Nature Communications, 8(1), Article 1. 10.1038/ncomms14435

59. Simberloff, D., Martin, J.-L., Genovesi, P., Maris, V., Wardle, D. A., Aronson, J., Courchamp, F., Galil, B., García-Berthou, E., Pascal, M., Pyšek, P., Sousa, R., Tabacchi, E., & Vilà, M. (2013). Impacts of biological invasions: What’s what and the way forward. Trends in Ecology & Evolution, 28(1), 58–66. 10.1016/j.tree.2012.07.013

60. Sittaro, F., Hutengs, C., & Vohland, M. (2023). Which factors determine the invasion of plant species? Machine learning based habitat modelling integrating environmental factors and climate scenarios. International Journal of Applied Earth Observation and Geoinformation, 116, 103158. 10.1016/j.jag.2022.103158

61. Smith, M. C., Pratt, P. D., & Rayamahji, M. B. (2022). Crown area predicts total biomass for Rhodomyrtus tomentosa, an invasive shrub in Florida. Invasive Plant Science and Management, 15(1), 61–66. 10.1017/inp.2022.8

62. Syphard, A. D., & Franklin, J. (2009). Differences in spatial predictions among species distribution modeling methods vary with species traits and environmental predictors. Ecography, 32(6), 907–918. 10.1111/j.1600-0587.2009.05883.x

63. Thuiller, W., Guéguen, M., Renaud, J., Karger, D. N., & Zimmermann, N. E. (2019). Uncertainty in ensembles of global biodiversity scenarios. Nature Communications, 10(1), Article 1. 10.1038/s41467-019-09519-w

64. van Kleunen, M., Dawson, W., Essl, F., Pergl, J., Winter, M., Weber, E., Kreft, H., Weigelt, P., Kartesz, J., Nishino, M., Antonova, L. A., Barcelona, J. F., Cabezas, F. J., Cárdenas, D., Cárdenas-Toro, J., Castaño, N., Chacón, E., Chatelain, C., Ebel, A. L., … Pyšek, P. (2015). Global exchange and accumulation of non-native plants. Nature, 525(7567), Article 7567. 10.1038/nature14910

65. Weigelt, P., König, C., & Kreft, H. (2020). GIFT – A Global Inventory of Floras and Traits for macroecology and biogeography. Journal of Biogeography, 47(1), 16–43. 10.1111/jbi.13623

66. Wisz, M. S., Hijmans, R. J., Li, J., Peterson, A. T., Graham, C. H., Guisan, A., & Group, N. P. S. D. W. (2008). Effects of sample size on the performance of species distribution models. Diversity and Distributions, 14(5), 763–773. 10.1111/j.1472-4642.2008.00482.x

67. Wohlwend, M. R., Craven, D., Weigelt, P., Seebens, H., Winter, M., Kreft, H., Zurell, D., Sarmento Cabral, J., Essl, F., van Kleunen, M., Pergl, J., Pyšek, P., & Knight, T. M. (2021). Anthropogenic and environmental drivers shape diversity of naturalized plants across the Pacific. Diversity and Distributions, 27(6), 1120–1133. 10.1111/ddi.13260

68. Wood, S. N., N., Pya, & Safken, B. (2016). Smoothing parameter and model selection for general smooth models (with discussion). Journal of the American Statistical Association, 111, 1548–1575.

69. Wüest, R. O., Zimmermann, N. E., Zurell, D., Alexander, J. M., Fritz, S. A., Hof, C., Kreft, H., Normand, S., Cabral, J. S., Szekely, E., Thuiller, W., Wikelski, M., & Karger, D. N. (2020). Macroecology in the age of Big Data – Where to go from here? Journal of Biogeography, 47(1), 1–12. 10.1111/jbi.13633

70. Ziller, S. R., de Sá Dechoum, M., Silveira, R. A. D., da Rosa, H. M., Motta, M. S., da Silva, L. F., Oliveira, B. C. M., & Zenni, R. D. (2020). A priority-setting scheme for the management of invasive non-native species in protected areas. NeoBiota, 62, 591–606.

71. Zizka, A., Silvestro, D., Andermann, T., Azevedo, J., Duarte Ritter, C., Edler, D., Farooq, H., Herdean, A., Ariza, M., Scharn, R., Svantesson, S., Wengström, N., Zizka, V., & Antonelli, A. (2019). CoordinateCleaner: Standardized cleaning of occurrence records from biological collection databases. Methods in Ecology and Evolution, 10(5), 744–751. 10.1111/2041-210X.13152

72. Zurell, D., Franklin, J., König, C., Bouchet, P. J., Dormann, C. F., Elith, J., Fandos, G., Feng, X., Guillera-Arroita, G., Guisan, A., Lahoz-Monfort, J. J., Leitão, P. J., Park, D. S., Peterson, A. T., Rapacciuolo, G., Schmatz, D. R., Schröder, B., Serra-Diaz, J. M., Thuiller, W., … Merow, C. (2020). A standard protocol for reporting species distribution models. Ecography, 43(9), 1261–1277. 10.1111/ecog.04960

73. Zurell, D., Zimmermann, N. E., Gross, H., Baltensweiler, A., Sattler, T., & Wüest, R. O. (2020). Testing species assemblage predictions from stacked and joint species distribution models. Journal of Biogeography, 47(1), 101–113. 10.1111/jbi.13608

