## Supplementary Material A for "Uncertainty in blacklisting potential Pacific plant invaders using species distribution models"

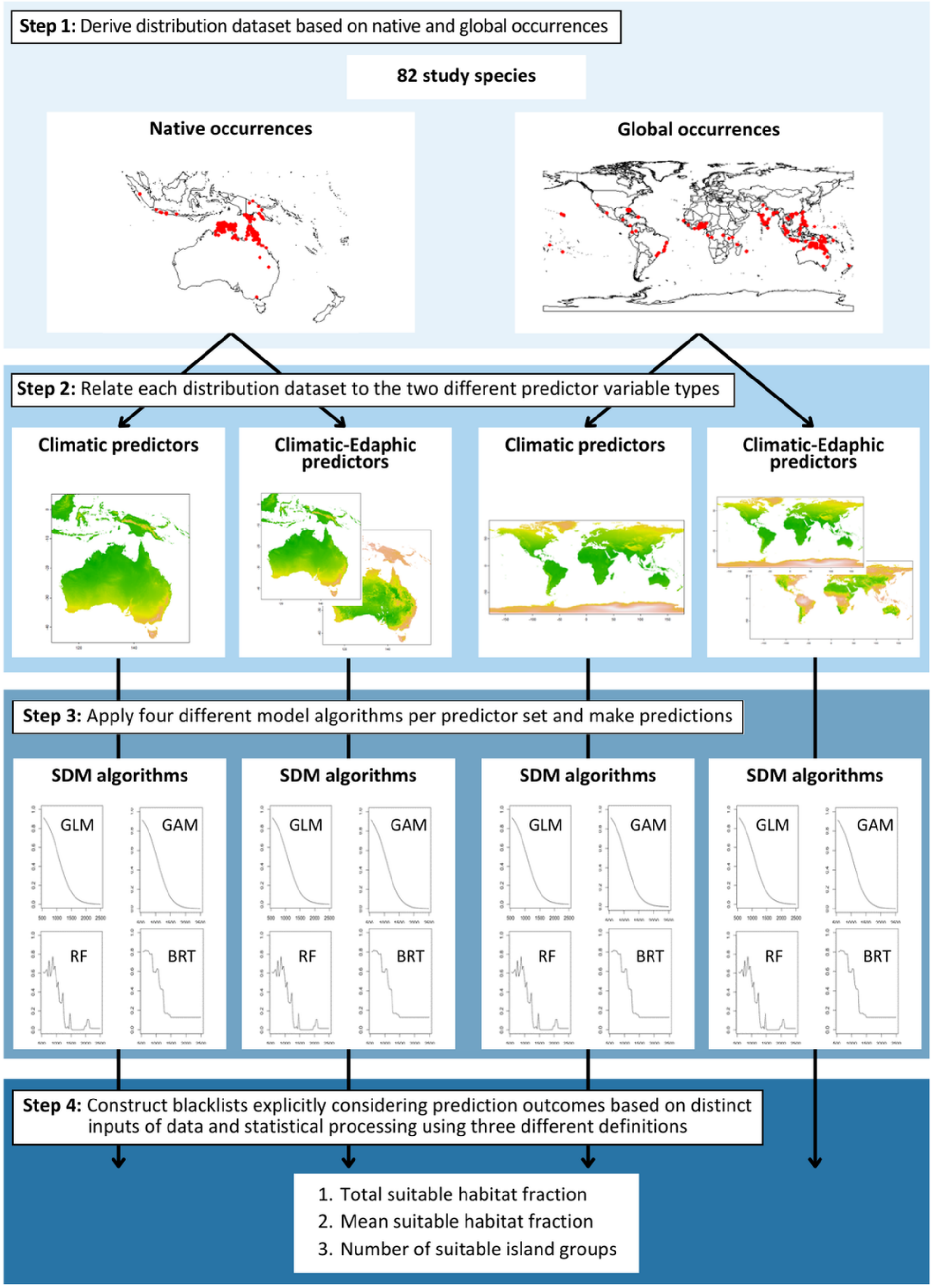


**Figure S1: Schematic representation of the workflow.** For all study species (n = 82), SDMs were fitted on opportunistic species data representative of their native or global niches, using two different predictor variable sets (climatic and edapho-climatic) and four different SDM algorithms. From the predicted habitat suitability maps, we derived blacklists using three distinct definitions. These blacklists were then used to quantify the uncertainty in ranking related to three different sources: species data, predictor type, and algorithm.

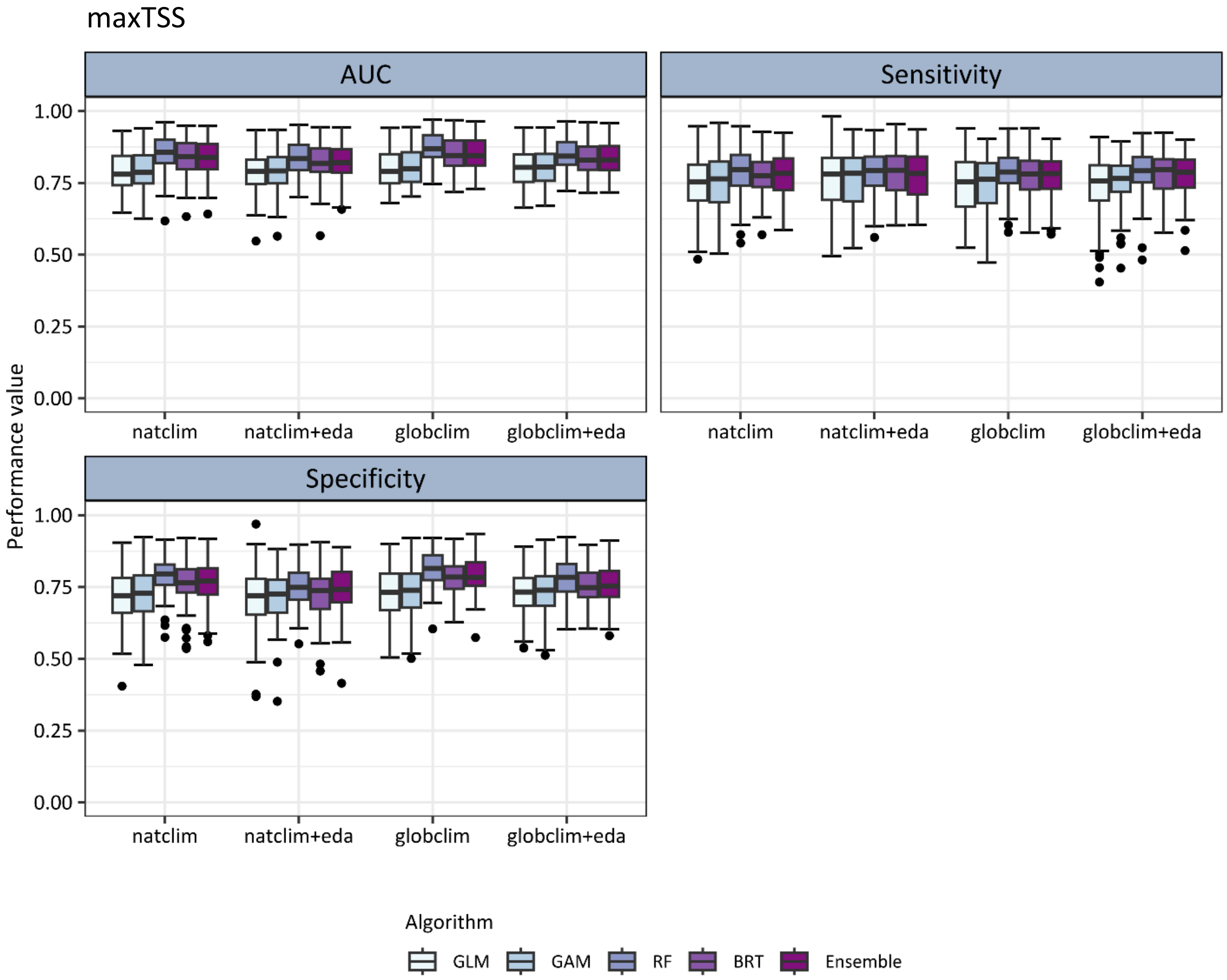


**Figure S2: Model validation in terms of AUC (Area Under the Receiver Operating Characteristic Curve), Sensitivity and Specificity.** For all studied non-native plant species (n = 82), the range of performance measures for the four applied model algorithms and the averaged ensemble model along the four different input datasets are displayed as boxplots. Models were validated using 5-fold cross-validation, and threshold-dependent performance measures were calculated using the threshold that maximises the True Skill Statistic (maxTSS). (GLM: generalised linear model; GAM: generalised additive model; RF: random forest; BRT: boosted regression tree; natclim: models based on native occurrences and climate data; globclim: models based on global occurrences and climate data; eda: models additionally considering edaphic data)

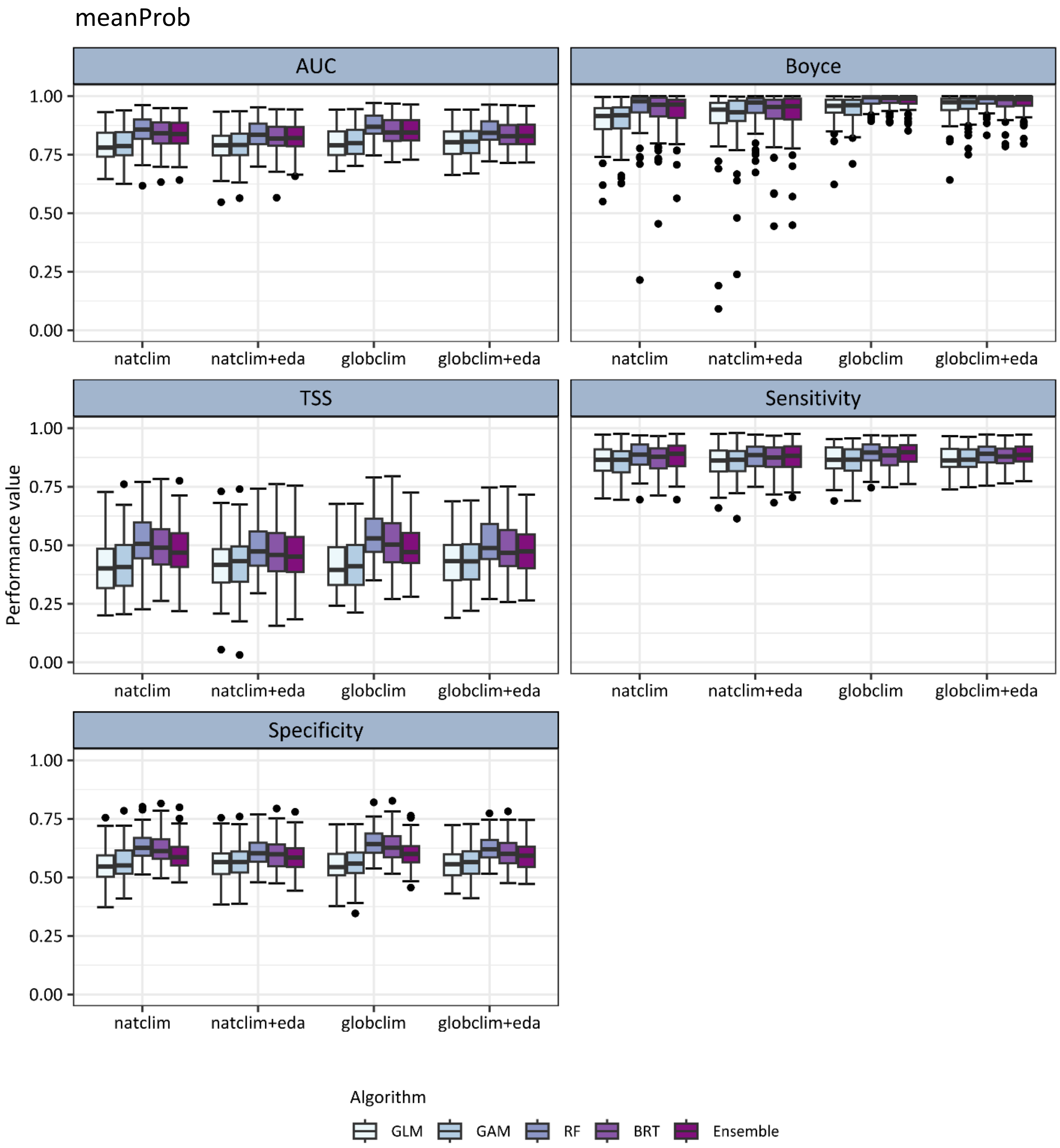


**Figure S3: Model validation in terms of AUC (Area Under the Receiver Operating Characteristic Curve), Boyce index, TSS (True Skill Statistic), Sensitivity and Specificity.** For all studied non-native plant species (n = 82), the range of performance measures for the four applied model algorithms and the averaged ensemble model along the four different input datasets are displayed as boxplots. Models were validated using 5-fold cross-validation, and threshold-dependent performance measures were calculated using the mean probability of training locations (meanProb) as the threshold. (GLM: generalised linear model; GAM: generalised additive model; RF: random forest; BRT: boosted regression tree; natclim: models based on native occurrences and climate data; globclim: models based on global occurrences and climate data; eda: models additionally considering edaphic data)

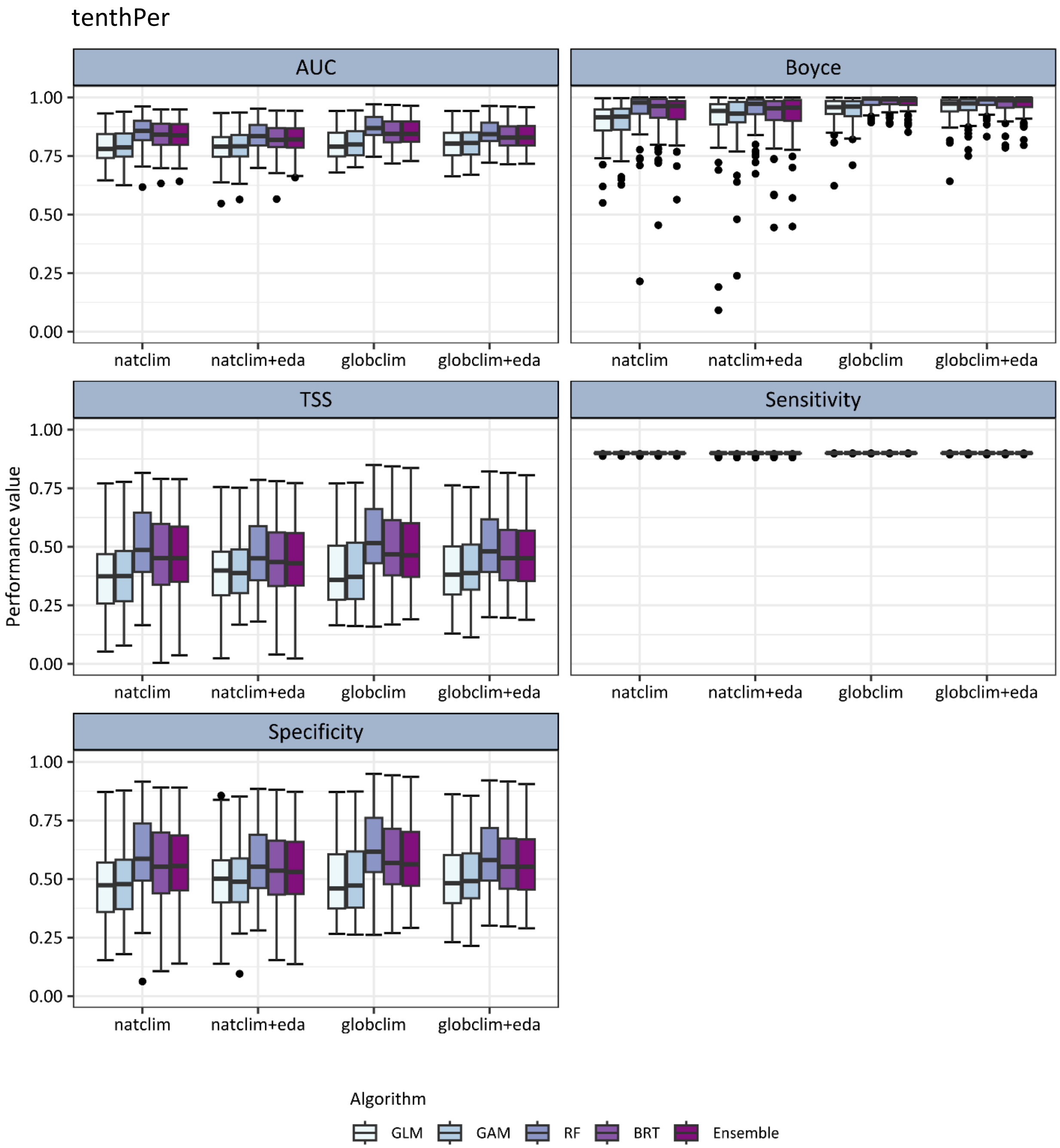


**Figure S4: Model validation in terms of AUC (Area Under the Receiver Operating Characteristic Curve), Boyce index, TSS (True Skill Statistic), Sensitivity and Specificity.** For all studied non-native plant species (n = 82), the range of performance measures for the four applied model algorithms and the averaged ensemble model along the four different input datasets are displayed as boxplots. Models were validated using 5-fold cross-validation, and threshold-dependent performance measures were calculated using the 10th percentile of probabilities at presence locations (tenthPer) as the threshold. (GLM: generalised linear model; GAM: generalised additive model; RF: random forest; BRT: boosted regression tree; natclim: models based on native occurrences and climate data; globclim: models based on global occurrences and climate data; eda: models additionally considering edaphic data)

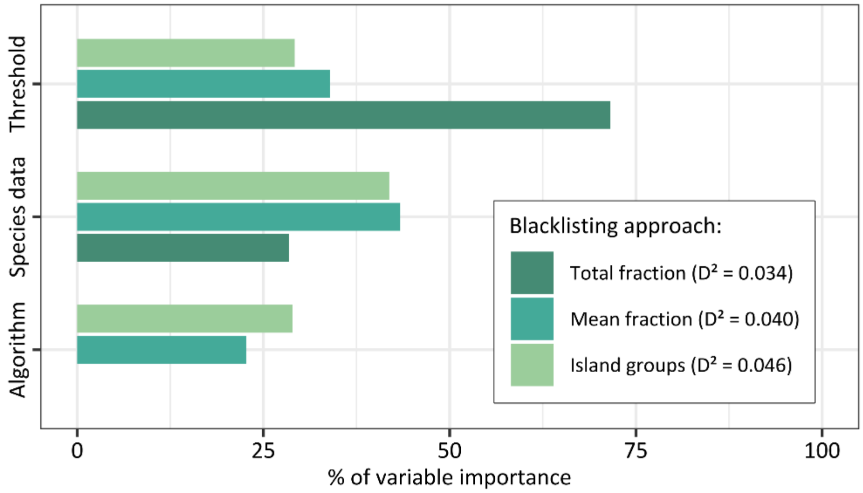


**Figure S5: Quantification of uncertainty sources in establishment risk scores underlying blacklist rankings, based on analysis of 49 Pacific island groups.** The bars display the influence in the choice of three different sources of uncertainty on variation in establishment risk scores: the algorithm (GLM, GAM, RF, BRT), the species data (global vs. native occurrences), and the thresholding method (maxTSS, meanProb, tenthPer). Variable importance was estimated using a random forest model that relates differences in establishment risk scores to the uncertainty factors. The values of variable importance per blacklisting approach were standardised to sum up to 100 %. The explained deviance $D^2$ of the models is noted in the legend. The models include establishment risk scores of all studied non-native plant species (n = 82) based on predictions to 49 Pacific island groups.

**Table S1: Quantification of uncertainty sources in blacklist rankings based on analysis of 49 island groups.** For each blacklist approach, the table presents the mean differences in species rankings, along with the proportion of cases showing changes or no changes in rank groupings. Ranks were grouped into priority bins of 10 to assess grouping changes (i.e., Top 10, Top 20, Top 30, etc.). The analysis considers the three different uncertainty sources: the SDM algorithm (GLM, GAM, RF, BRT), the species data (global vs. native occurrences), and the thresholding method (maxTSS, meanProb, tenthPer). Results are based on blacklisting assessments of all studied non-native plant species (n = 82), predicted across 49 Pacific island groups covered by climatic data.

| Blacklist | Factor | Mean ranking differences | No change in grouping (%) | Change in grouping (%) |
|---|---|---|---|---|
| **Total fraction** | Algorithm | 15.24 | 20 | 80 |
| | Species data | 12.25 | 34 | 66 |
| | Threshold | 8.07 | 41 | 59 |
| **Mean fraction** | Algorithm | 16.17 | 19 | 81 |
| | Species data | 12.48 | 33 | 67 |
| | Threshold | 8.00 | 41 | 59 |
| **Island groups** | Algorithm | 7.51 | 55 | 45 |
| | Species data | 5.80 | 64 | 36 |
| | Threshold | 4.59 | 68 | 32 |

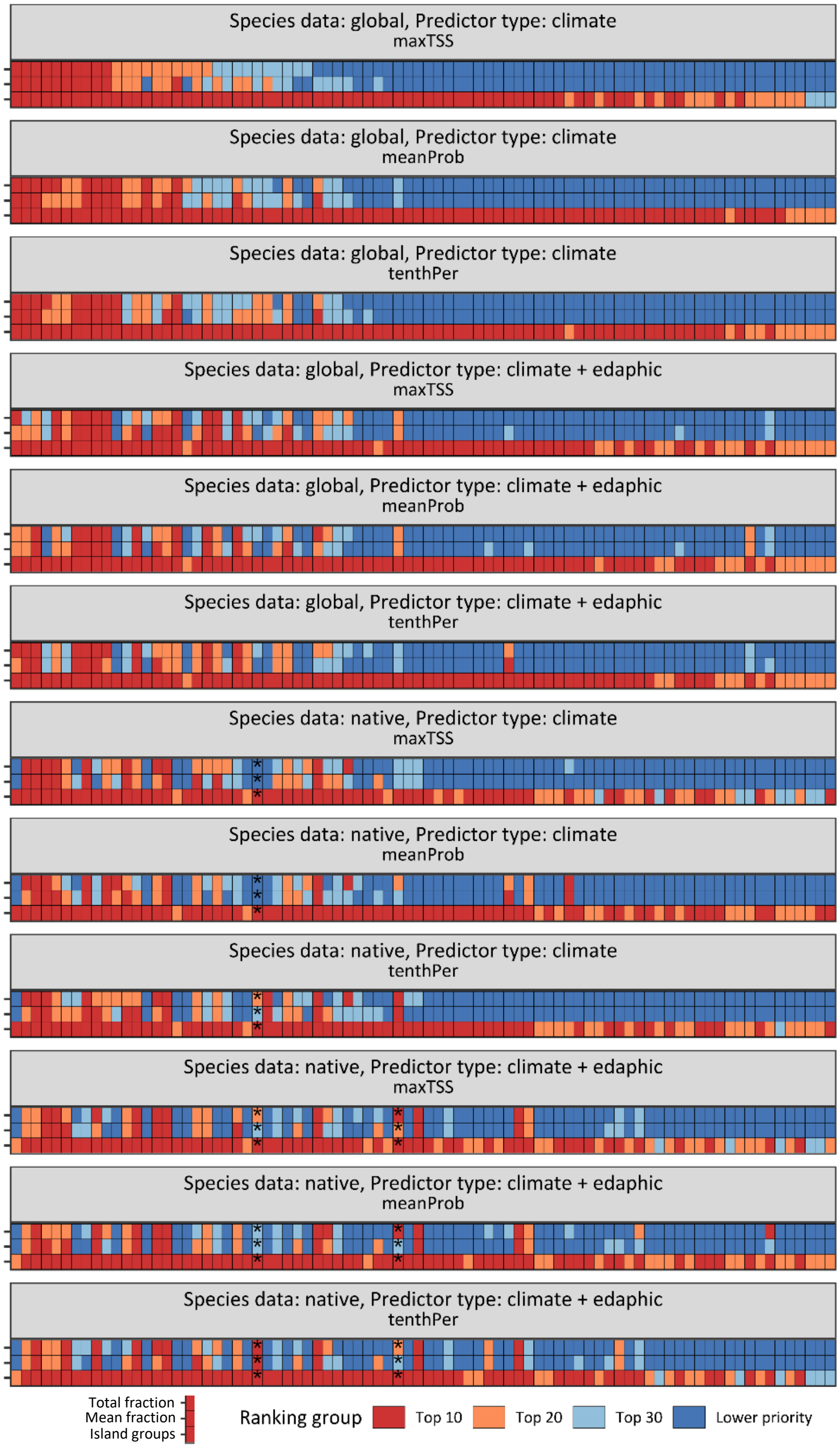


**Figure S6: Final blacklist groupings based on analysis of 25 Pacific island groups.** The final blacklist groupings of all three blacklisting approaches over four different input datasets are displayed. The groupings are illustrated for each of the three thresholding methods (maxTSS, meanProb, tenthPer), based on the predictions of ensemble models to 25 Pacific island groups covered by climatic and edaphic data. Every non-native plant species (n = 82) is depicted along a vertical line of cells. The horizontal lines of cells per predictor set represent the three blacklisting approaches, with cell colour indicating the rank grouping for each species. Species ranks were categorised into four rank groups: Top 10, Top 20, Top 30, and Lower priority. Predictive performance values of ensemble models below or equal to a Boyce index of 0.7 are marked with an asterisk.

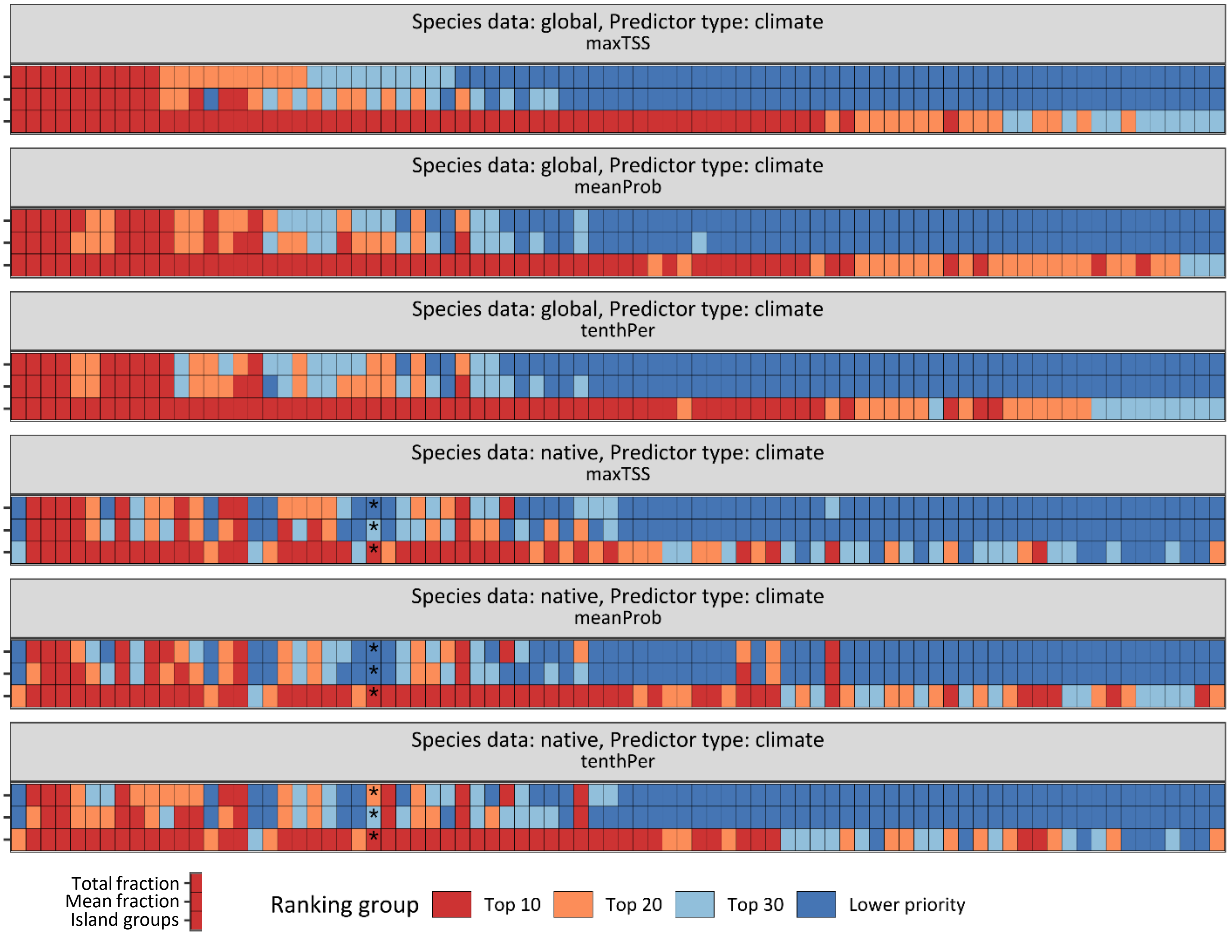


**Figure S7: Final blacklist groupings based on analysis of 49 Pacific island groups.** The final blacklist groupings of all three blacklisting approaches over two different input datasets are displayed. The groupings are illustrated for each of the three thresholding methods (maxTSS, meanProb, tenthPer), based on the predictions of ensemble models to 49 Pacific island groups solely covered by climatic data. Every non-native plant species (n = 82) is depicted along a vertical line of cells. The horizontal lines of cells per predictor set represent the three blacklisting approaches, with cell colour indicating the rank grouping for each species. Species ranks were categorised into four rank groups: Top 10, Top 20, Top 30, and Lower priority. Predictive performance values of ensemble models below or equal to a Boyce index of 0.7 are marked with an asterisk.

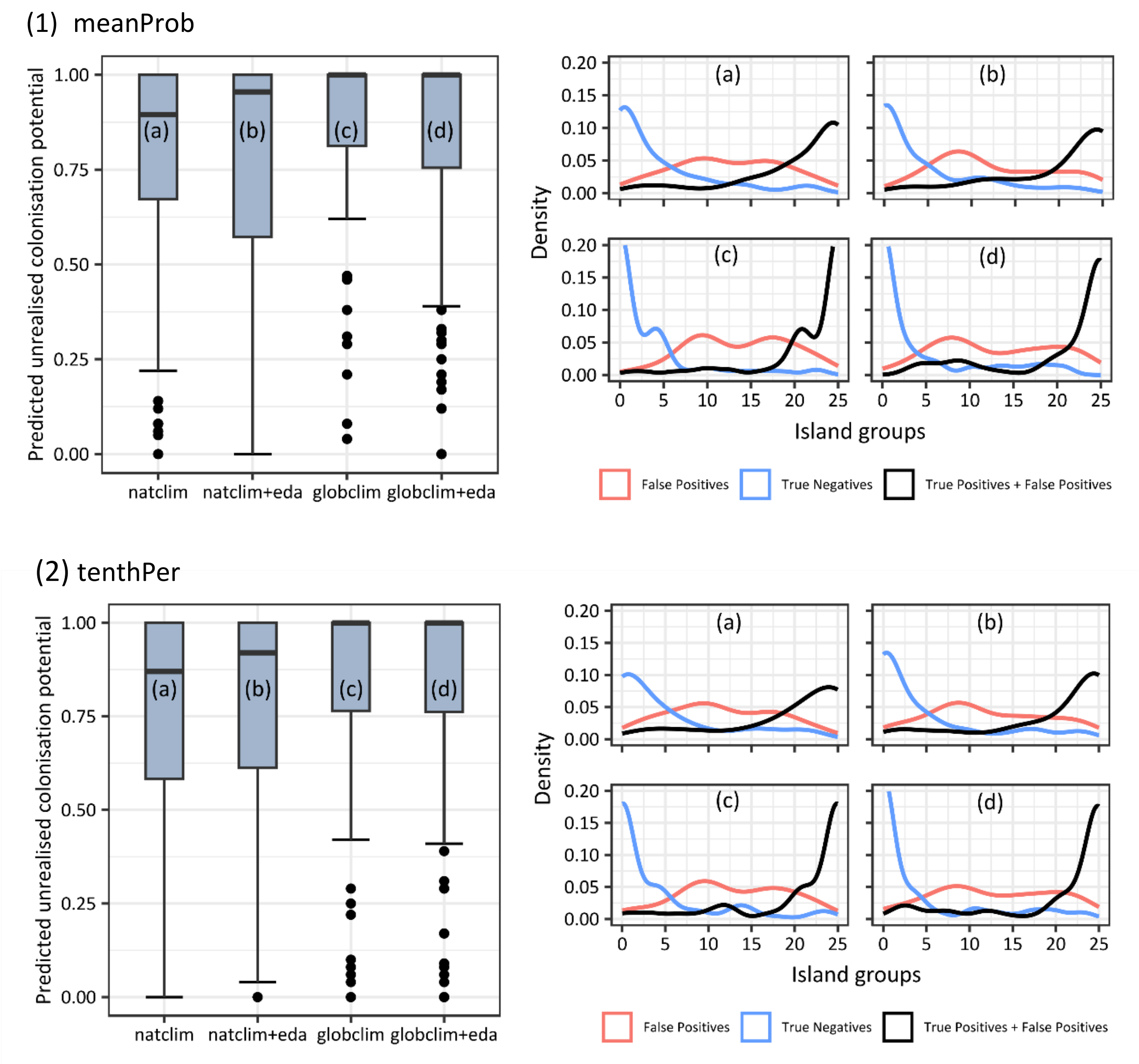


**Figure S8: Predicted Pacific-wide unrealised colonisation potential based on analysis of 25 Pacific island groups.** The figure consists of two parts: an upper panel (1) and a lower panel (2), each comprising a left and right subpanel. In both parts, the left subpanels display boxplots of the predicted unrealised colonisation potential for all studied non-native plant species (n = 82), based on the ensemble predictions of 25 Pacific island groups covered by edaphic and climatic data along all four input datasets (a-d; natclim: models based on native occurrences and climate data; globclim: models based on global occurrences and climate data; eda: models additionally considering edaphic data). The predicted unrealised colonisation potential per species represents the False Positive Rate defined as the number of island groups that are hitherto unoccupied by the non-native species but predicted as suitable, divided by the total number of island groups where the species is currently absent. The right subpanels show the density distribution of island groups with false positives (predicted but absent; red line), true negatives (blue line), and all predicted presences (black line), depicted for all study species using the same ensemble predictions (a-d). Thresholding methods differed between the two parts: in the upper panel (1), predictions were binarised using the mean of predicted probabilities at training locations (meanProb); in the lower panel (2), the 10th percentile of predicted probabilities at presence locations (tenthPer) was used as the threshold.

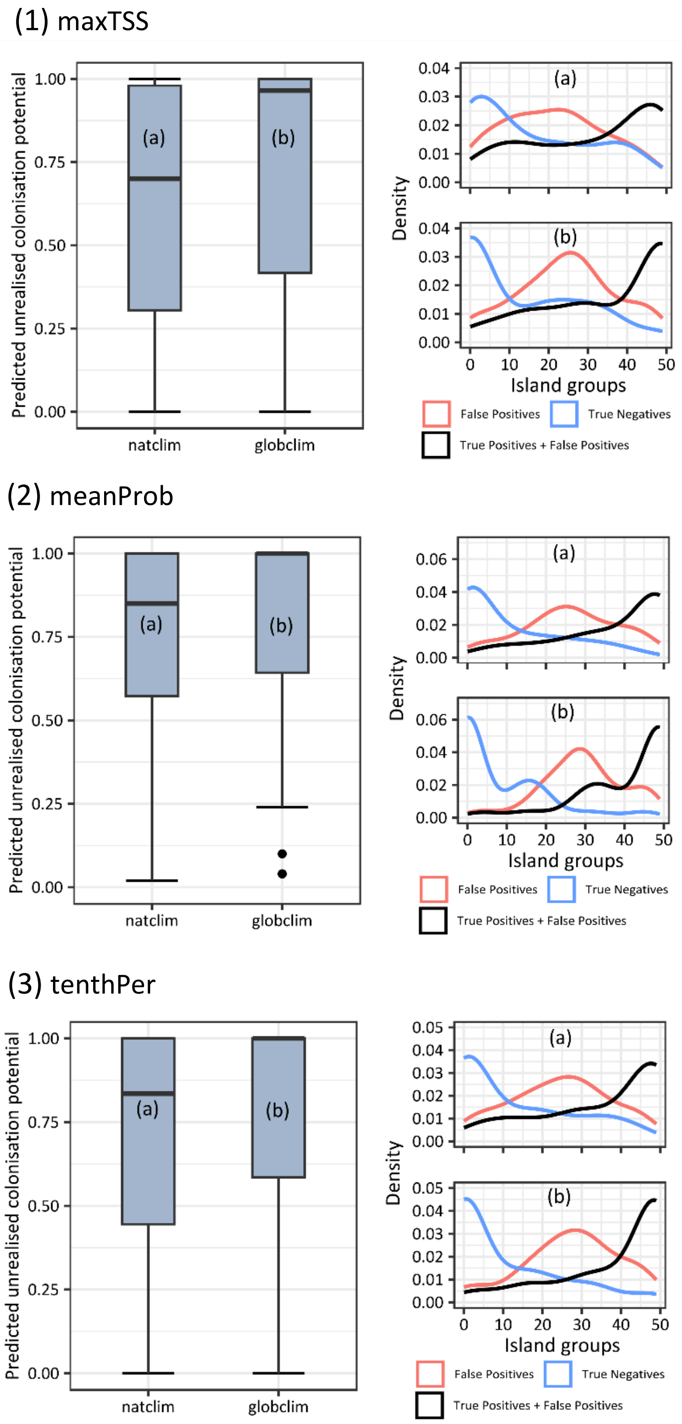


**Figure S9: Predicted Pacific-wide unrealised colonisation potential based on analysis of 49 Pacific island groups.** The figure consists of three parts: an upper panel (1), a middle panel (3), and a lower panel (2), each comprising a left and right subpanel. On the left, the predicted Pacific-wide unrealised colonisation potential for all studied non-native plant species (n = 82) is displayed as boxplots, based on the ensemble predictions of 49 Pacific island along the two predictor sets solely including climate data (a-b). The predicted unrealised colonisation potential per species represents the False Positive Rate defined as the number of island groups that are hitherto unoccupied by the non-native species but predicted as suitable divided by the total number of island groups where the species is currently absent. On the right, the density of island groups with false positives (predicted but absent; red line), true negatives (blue line), and all predicted presences (black line) is depicted for all study species using the same ensemble predictions (a-b). (natclim: models based on native occurrences and climate data; globclim: models based on global occurrences and climate data). Thresholding methods differed between the three parts: in the upper panel (1), predictions were binarised using the threshold that maximises the True Skill Statistic (maxTSS); in the middle panel (2), predictions were binarised using the mean of predicted probabilities at training locations (meanProb); in the lower panel (3), the 10th percentile of predicted probabilities at presence locations (tenthPer) was used as the threshold.

**Data Acknowledgements**

We acknowledge the herbaria that contributed data to this work: U, MBML, UFRN, ARIZ, BRIT, HUAL, M, CSLA, CMC, CM, BRNU, BIRM, BHSC, BERN, BASBG, ABD, MAL, MACB, LU, LOB, LMU, LAU, KSP, KR, KATH, KAND, IBF, HSU, HLU, GSW, GJO, FSC, ERE, DES, DEE, TUCH, TOGO, TBI, TAA, SNUA, SJC, SHIN, RENO, PI, NMW, NMLU, NHG, NEBC, NCCE, MSUB, YRK, YA, WSCO, WCUH, WB, VPI, VALPL, UNSW, A, AAS, AAU, ABH, ACAD, AD, AK, AKPM, ALTA, AMNH, ARAN, ASU, B, BABY, BAS, BC, BCN, BEREA, BG, BM, BOON, BOUM, BR, BRI, C, CAN, CANB, CAS, CBM, CDBI, CHAS, CHR, CHRB, CHSC, CICY, CNS, COA, COFC, COI, COL, COLO, CONC, CORD, CP, CU, CVRD, DAO, DAV, DBG, DNA, E, EA, EKY, EMMA, ER, F, FAU, FCO, FLAS, FR, FRU, FTG, FULD, FURB, G, GAT, GB, GH, GLM, GMNHJ, GZU, HAL, HAST, HIB, HNT, HO, HSS, HU, HUJ, HYO, IAC, IBK, IBSC, ICEL, ICESI, ICN, INEGI, INM, INPA, IPA, ISKW, JBAG, JBGP, JCT, KYO, JYV, K, KMN, KOR, KPM, KSTC, KTU, KU, KUN, LAGU, LBG, LD, LE, LEB, LSU, LTB, LTR, MA, MACF, MAF, MAK, MB, MBK, MBM, MEL, MELU, MFU, MGC, MICH, MIN, MMMN, MNHM, MNHN, MO, MSB, MSC, MT, MTMG, MUB, NAC, NAS, NCSC, NCU, NE, NH, NHM, NHT, NMNL, NMR, NMSU, NSPM, NSW, NU, NY, O, OBI, OSA, OSC, P, PAMP, PE, PERTH, POM, QFA, QUE, REG, RELC, RNG, RSA, S, SALA, SAN, SANT, SAPS, SASK, SBBG, SCFS, SD, SDSU, SFV, SJSU, STU, SVG, TAI, TAIF, TALL, TAM, TAMU, TAN, TASH, TEF, TENN, TEX, TI, TKPM, TNS, TOYA, TRH, TROM, TRT, TRTE, UADY, UAM, UBC, UBT, UCR, UCS, UCSB, UCSC, UESC, UFG, UFMA, UFMT, UFS, UJAT, ULM, ULS, UME, UNB, UNM, UPNA, UPS, US, USCH, USF, USP, UT, UTEP, UWO, V, VAL, VIT, VT, W, WAG, WAT, WII, WIN, WOLL, WU, YAMA, Z, ZT, accessed via the BIEN dataset.
