## Supplementary Material C for "Uncertainty in blacklisting potential Pacific plant invaders using species distribution models"

### – ODMAP Protocol –

Valén Holle, Anna Rönnfeldt, Katrin Schifferle, Juliano Sarmento Cabral, Dylan Craven, Tiffany Knight, Hanno Seebens, Patrick Weigelt, Damaris Zurell

2025-06-10

---

### Overview

#### *Authorship*

Contact:

Study link:

#### *Model objective*

Model objective: Mapping and interpolation

Target output: suitable vs. unsuitable habitat

#### *Focal Taxon*

Focal Taxon: Vascular plant species

#### *Location*

Location: Pacific region (up to 49 island groups)

#### *Scale of Analysis*

Spatial extent: 130, -75, -40, 40 (xmin, xmax, ymin, ymax)

Spatial resolution: 1 km

Temporal extent: 1900-2023

Temporal resolution: one studied time period (1900-2023)

Boundary: political, natural

#### *Biodiversity data*

Observation type: citizen science

Response data type: presence-only

*Predictors*

Predictor types: climatic, edaphic

*Hypotheses*

Hypotheses: The species distributions are mainly determined by climate and soil.

*Assumptions*

Model assumptions: We assumed that climate and soil are the key explanatory variables for species distributions, species are in equilibrium with environment, observation biases are insignificant.

*Algorithms*

Modelling techniques: glm, gam, randomForest, brt

Model complexity: We used an intermediate level of model complexity to prevent overfitting for species with smaller sample sizes and to ensure that predictions to new places remain robust.

Model averaging: We used the arithmetic mean to combine all four model algorithms into one ensemble.

*Workflow*

Model workflow: We downloaded species data from GBIF and BIEN, cleaned the coordinates, and determined the biogeographical status of plant occurrences as either native or introduced based on three different sources: WCVP, GIFT, and GIoNAF. We generated background data by randomly selecting points within a buffer distance of 200 km from the occurrence locations, using a presence-absence ratio of 1:10. To avoid spatial autocorrelation, we then thinned all presences and pseudo-absences using a 3 km threshold. We obtained climate data (CHELSA) and edaphic data (SoilGrids) at a 1 km resolution. Given that our study aimed to quantify the uncertainty in species input data and environmental input data for SDM-based blacklists, we intersected the two types of species data (native vs. global) with the two types of environmental data (climatic vs. edapho-climatic), resulting in four combinations of input data for each species. For each of these four SDM input datasets, we selected the four most important and weakly correlated predictor variables for model construction. SDMs were fitted using four statistical algorithms (GLM, GAM, RF, BRT). For the regression-based methods (GLM, GAM), we used a presence-absence ratio of 1:10 but applied equal weighting during model building. For the machine-learning methods, we used ten replicate sets of background data in a presence-absence ratio of 1:1, generating 10 different models, which were then averaged. Additionally, model ensembles were constructed using the arithmetic mean of predictions over all four model algorithms. The SDM performance was evaluated using a 5-fold cross-validation by computing AUC, TSS, sensitivity, specificity, and the Boyce index. Based on each model and their ensemble, we predicted potential presence and absence across the Pacific using three different thresholds (maxTSS, meanProb, tenthPer). Based on the quantified area of predicted suitable habitat in km$^2$, we separately constructed a blacklist for each combination of species input data, environmental input data and SDM algorithm, as well as a final blacklist with ensembles of SDM algorithms.

#### *Software*

Software: R 4.3.1 with [1] BIEN_1.26 [2] conflicted_1.2.0.9000 [3] CoordinateCleaner_3.0.1 [4] dismo_1.3-14 [5] doParallel_1.0.17 [6] ecospat_4.0.0 [7] fasterize_1.0.4 [8] foreach_1.5.2 [9] GIFT_1.0.0 [10] gbm_2.1.8.1 [11] ggplot2_3.5.1 [12] ggtext_0.1.2 [13] lcvplants_2.1.0 [14] maps_3.4.1 [15] mgcv_1.8-42 [16] PresenceAbsence_1.1.11 [17] randomForest_4.7-1.1 [18] readr_2.1.4 [19] rgbif_3.7.5 [20] rWCVP_1.2.4 [21] sf_1.0-16 [22] sfheaders_0.4.3 [23] showtext_0.9-7 [24] taxize_0.9.100 [25] terra_1.7-55 [26] tibble_3.2.1
[27] tidyverse_2.0.0 [28] units_0.8-5 [29] viridis_0.6.4

Code availability: All scripts are available in <>.

Data availability: All data are publicly available and can be downloaded using the available scripts.

### Data

#### *Biodiversity data*

Taxon names: *Acacia auriculiformis, Acanthospermum australe, Alternathera brasiliana, Alysicarpus vaginalis, Anemone hupehensis, Anredera cordifolia, Arrhenatherum elatius, Asclepis curassavica, Bellis perennis, Botriochloa bladhii, Brachiara subquadripara, Brassica rapa, Breynia disticha, Caesalpinia decapetala, Calotropis gigantea, Capsicum frutescens, Cereus hildmannianus, Chloris radiata, Cirsium vulgare, Crassocephalum crepidioides, Crotalaria juncea, Crotalaria pallida, Delonix regia, Desmanthus pernambucanus, Dieffenbachia seguine, Digitaria eriantha, Digitaria horizontalis, Echinochloa esculenta, Emilia sonchifolia, Erigeron bonariensis, Eugenia uniflora, Euphorbia tirucalli, Ficus religiosa, Gamochaeta americana, Gnaphalium pensylvanicum, Gnaphalium purpureum, Grevillea banksii, Grevillea robusta, Haematoxylum campechianum, Justicia spicigera, Lablab purpureus, Lamarckia aurea, Laportea aestuans, Lespedeza cuneata, Lolium arundinaceum, Lophostemon confertus, Olea europea, Paraserianthes lophantha, Passiflora caerulea, Pentas lanceolata, Phleum pratense, Phoenix dactylifera, Physalis angulata, Picris hieracioides, Pistia stratiotes, Portulaca pilosa, Prunus persica, Psidium guajava, Rhodomyrtus tomentosa, Rubus ellipticus, Rumex spinosus, Schefflera arboricola, Senna pendula, Sigesbeckia orientalis, Sphagneticola trilobata, Stachys arvensis, Stylosanthes guianensis, Syzygium malaccense, Tagetes minuta, Tanacetum parthenium, Tecoma stans, Tillandsia usneoides, Tithonia diversifolia, Tragus berteronianus, Tribulus terrestris, Tridax procumbens, Vanilla planifolia, Verbena bonariensis, Verbena rigida, Verbesina encelioides, Vitis vinifera, Xanthium strumarium*

Taxonomic reference system: GBIF/BIEN backbone taxonomy

Ecological level: species

Data sources: GBIF and BIEN (accessed in June 2023)

Sampling design: opportunistic

Sample size: For models based on native occurrences, we used a minimum number of presences n = 40. Final presences for the native models ranged from 42 to 39,486. For models based on global occurrences, we used a minimum number of n = 40 native presences and n = 40 non-native presences. Final presences for the global model ranged from 122 to 70,495.

Scaling: We spatially thinned the data to 3 km.

Cleaning: We removed duplicate observations; observations with missing coordinates, equal coordinates, coordinate uncertainty > 10 km; coordinates within a 1,000 m radius of country centroids, within a 10,000 m radius of capitals, within a 100 m radius of biodiversity institutions, within 1° radius of GBIF headquarters in Copenhagen.

Absence data: N/A

Background data: We generated background data within a 200 km buffer around presence locations while excluding those. We derived 10 times as many background points as presence points.

#### *Data partitioning*

Training data: GLMs and GAMs were fitted based on all data points. For RFs and BRTs, 10 replicate models in a presence-absence ratio of 1:1 were fitted.

Validation data: We used a 5-fold cross-validation.

#### *Predictor variables*

Predictor variables: 15 bioclimatic variables (bio1, bio2, bio3, bio4, bio5, bio6, bio7, bio10, bio11, bio12, bio13, bio14, bio15, bio16, bio17); 14 edaphic variables (soil organic carbon content, total nitrogen, coarse fragments, pH, cation exchange capacity, bulk density, sand content, clay content, silt content, soil water content at three different pressure heads)

Data sources: Chelsa Version 2.1; SoilGrids Version 2.0

Spatial extent: global

Spatial resolution: 1 km

Coordinate reference system: WGS 84 (EPSG:4326)

Temporal extent: Chelsa: 1981-2010; SoilGrids: data collected up until 2017

Data processing: We averaged the soil data over the three downloaded depth intervals (30 cm).

Dimension reduction: We excluded bio8, bio8, bio18, and bio19 as these are variables that combine temperature and precipitation data. We pursued to avoid artefacts leading to discontinuities in interpolated surfaces.

### Model

#### *Multicollinearity*

Multicollinearity: We checked for multi-collinearity and only retained the more important variable from highly correlated pairs with Spearman correlations $|r| > 0.7$. Variable importance was assessed using the Boyce index, determined based on univariate GLMs and a 5-fold cross-validation. We considered the four variables that were ranked most important in the list of weakly correlated predictors for modelling.

#### *Model settings*

glm: family (binomial), formula (linear and quadratic terms), weights (equal weights of presences and absences)

gam: family (binomial), smoothTerms (non-parametric cubic smoothing splines (k =4))

randomForest: ntree (1000), maxnodes (20)

brt: distribution (Bernoulli), interactionDepth (2), shrinkage (optimised learning rate to yield tree numbers between 1,000 and 5,000), bagFraction (0.75)

#### *Model estimates*

Coefficients: N/A

#### *Model selection - model averaging - ensembles*

Model averaging: We used the arithmetic mean to average the predictions of the 10 replicate models of RFs and BRTs.

Model ensembles: We used the arithmetic mean to average the predictions of the four different algorithms.

#### *Analysis and Correction of non-independence*

Spatial autocorrelation: N/A

#### *Threshold selection*

Threshold selection: threshold that maximises the true skill statistic (maxTSS), the mean probability of training points (meanProb), and the $10^{th}$ percentile of probabilities at training presences (tenthPer)

### Assessment

#### *Performance statistics*

Performance on training data: N/A

Performance on validation data: AUC, True positive rate, True negative rate, TSS, Boyce

Performance on test data: N/A

#### *Plausibility check*

Response shapes: N/A

Expert judgement: N/A

### Prediction

#### *Prediction output*

Prediction unit: continuous habitat suitability; suitable vs. unsuitable habitat (binary measure)

#### *Uncertainty quantification*

Scenario uncertainty: N/A

Novel environments: N/A
